# Evaluation of Polygenic Prediction Methodology within a Reference-Standardized Framework

**DOI:** 10.1101/2020.07.28.224782

**Authors:** Oliver Pain, Kylie P. Glanville, Saskia P. Hagenaars, Saskia Selzam, Anna E. Fürtjes, Héléna A. Gaspar, Jonathan R. I. Coleman, Kaili Rimfeld, Gerome Breen, Robert Plomin, Lasse Folkersen, Cathryn M. Lewis

## Abstract

**Background:** The predictive utility of polygenic scores is increasing, and many polygenic scoring methods are available, but it is unclear which method performs best. This study evaluates the predictive utility of polygenic scoring methods within a reference-standardized framework, which uses a common set of variants and reference-based estimates of linkage disequilibrium and allele frequencies to construct scores.

**Methods:** Eight polygenic score methods were tested: p-value thresholding and clumping (pT+clump), SBLUP, lassosum, LDPred1, LDPred2, PRScs, DBSLMM and SBayesR, evaluating their performance to predict outcomes in UK Biobank and the Twins Early Development Study (TEDS). Strategies to identify optimal p-value threshold and shrinkage parameters were compared, including 10-fold cross validation, pseudovalidation and infinitesimal models (with no validation sample), and multi-polygenic score elastic net models.

**Results:** LDPred2, lassosum and PRScs performed strongly using 10-fold cross-validation to identify the most predictive p-value threshold or shrinkage parameter, giving a relative improvement of 16-18% over pT+clump in the correlation between observed and predicted outcome values. Using pseudovalidation, the best methods were PRScs and DBSLMM, with a relative improvement of >10% over other pseudovalidation and infinitesimal methods (lassosum, SBLUP, SBayesR, LDPred1, LDPred2). PRScs pseudovalidation was only 3% worse than the best polygenic score identified by 10-fold cross validation. Elastic net models containing polygenic scores based on a range of parameters consistently improved prediction over any single polygenic score.

**Conclusion:** Within a reference-standardized framework, the best polygenic prediction was achieved using LDPred2, lassosum and PRScs, modeling multiple polygenic scores derived using multiple parameters. This study will help researchers performing polygenic score studies to select the most powerful and predictive analysis methods.

## Introduction

In personalized medicine, medical care is tailored for the individual to provide improved disease prevention, prognosis, and treatment. Genetics is a potentially powerful tool for providing personalized medicine as genetic variation accounts for a large proportion of individual differences in health and disease [1]. Furthermore, an individual’s genetic sequence is stable across the lifespan, enabling predictions long before the onset of most diseases. Although genetic information is used to predict rare Mendelian genetic disorders, such as breast cancer based on *BRCA1/2* variants, our ability to predict common disorders using genetic information is currently insufficient for clinical implementation. This is due to the increased etiological complexity of common disorders, with complex interplay between genetic and environmental factors, and the highly polygenic genetic architecture with contributions from many genetic variants with small effect sizes [2]. However, genome-wide association studies (GWAS), used to detect common genetic associations, are rapidly increasing in sample size, and are identifying large numbers of novel and robust genetic associations for health-related outcomes [3]. This growing source of information is also improving our ability to predict an individual’s disease risk or measured trait based on their genetic variation [4, 5].

An individual’s genetic risk for an outcome can be summarized in a polygenic score, calculated from the number of trait-associated alleles carried. The contributing variants are typically weighted by the magnitude of effect they confer on the outcome of interest, estimated in a reference GWAS. There are several challenges in performing a well-powered polygenic score analysis. Firstly, GWAS effect-sizes are inflated through Winner’s curse, and unbiased estimates can only be obtained through an independent training sample, with these effect-size estimates then used to calculate polygenic scores in a further independent sample [6]. Secondly, to maximize polygenic prediction accuracy, the GWAS summary statistics must be adjusted to account for the linkage disequilibrium (LD) between genetic variants, to avoid double counting the non-independent effect of variants in high LD, and account for varying degrees of polygenicity across outcomes, i.e. the number of genetic variants affecting the outcome [6]. LD can be accounted for using LD-based clumping of GWAS summary statistics, removing variants in LD with lead variants within each locus, and polygenicity is accounted for by applying multiple GWAS *p*-value thresholds (pT) to select the effect alleles included in the polygenic score [4, 5]. This pT+clump approach is conceptually simple and computationally scalable [7]. However, using a hard LD threshold in clumping to retain or remove variants from the polygenic score calculation can potentially reduce the variance explained by the polygenic score. Alternative summary statistic-based polygenic score methods retain all genetic variants by modelling both the LD between variants and the polygenicity of the outcome [8–14]. These methods use estimates of LD to jointly estimate the effect of nearby genetic variation maximizing the signal captured, and generally apply a shrinkage parameter to the genetic effects to reduce overfitting and allow for varying degrees of polygenicity across outcomes.

Polygenic scoring methods can lead to overfitting of genetic effects due to the p-value based selection of variants or joint estimation of many genetic effects. To avoid this overfitting, genetic effect size estimates can be reduced using shrinkage methods to improve the generalizability of the model. Shrinkage methods for polygenic scoring can be separated into frequentist penalty-based methods (e.g. lasso regression-based lassosum [10], summary-based best linear unbiased prediction (SBLUP) [9]) and Bayesian methods that shrink estimates to fit a prior distribution of effect sizes, such as LDPred1 [8], LDPred2 [13], PRScs [11], SBayesR [12], and DBSLMM [14]. Each of these methods have been shown to improve the predictive utility of polygenic scores over those derived using the pT+clump approach. In comparisons between methods the findings are mixed: some studies have similar results across methods [15], while papers developing a new method often report that the developed method out-performs chosen other methods. To our knowledge no independent study has yet compared all approaches.

Five methods (pT+clump, LDPred1, LDPred2, lassosum and PRScs) generate multiple polygenic scores from user-defined tuning parameters. To determine which tuning parameter provides optimal prediction, the polygenic scores must first be tested in an independent ‘tuning’ sample. The pT+clump approach applies p-value thresholds to select variants included in the polygenic score, whereas LDPred1, LDPred2, lassosum and PRScs apply shrinkage parameters to adjust the GWAS effect sizes. In addition, lassosum, PRScs and LDPred2 provide a pseudovalidation approach, whereby a single optimal shrinkage parameter is estimated based on the GWAS summary statistics alone, and therefore do not require a tuning sample. SBayesR and DBSLMM can be considered pseudovalidation approaches as they also do not require a tuning sample to identify optimal parameters. Another approach to derive polygenic scores is to assume an infinitesimal model, as is done by SBLUP and the infinitesimal models of LDPred1 and 2 [16]. Similar to pseudovalidation approaches, no tuning sample is required when assuming an infinitesimal model. Rather than selecting a single tuning parameter, some studies have suggested that combining polygenic scores across p-value thresholds whilst taking into account their correlation using either PCA or model stacking can improve prediction [17, 18].

Polygenic scores are a useful research tool, as well as a promising potential tool for personalized healthcare through prediction of disease risk, prognosis, and treatment response [19]. However, polygenic scores calculated in a clinical setting should be valid for a single target sample and thus need to be constructed using a reference-standardized framework. Here, the polygenic score is independent of any properties specific to the target sample, including the genetic variation available, and the LD and minor allele frequency (MAF) estimates. In a reference-standardized approach, the genetic variants considered can be standardized by using only single nucleotide polymorphisms (SNPs) that are commonly available after imputation, such as variation within the HapMap3 reference [20]. The LD and MAF estimates can be standardized by using an ancestry matched individual-level genetic dataset such as 1000 Genomes [21]. Determining these properties (SNPs, LD, MAF) in reference data provides a practical approach for estimating polygenic scores for an individual, making them comparable to polygenic scores for other individuals of the same ancestry [22]. Use of a reference-standardized framework also offers advantages by improving the comparability of polygenic scores across cohorts. Several polygenic scoring methods now recommend the use of HapMap3 SNPs and precomputed external LD estimate references [11–13], in line with a reference-standardized approach.

In this study, we perform an extensive comparison of polygenic scoring methods within a reference-standardized framework. We evaluate the predictive utility of models for outcomes in UK Biobank and TEDS, combining information across tuning parameters. We evaluate eight polygenic scoring methods and apply different modelling strategies to select optimal tuning parameters to establish the combinations that perform consistently well. The reference-standardized framework increases the generalizability of results and provides a resource for future studies investigating polygenic prediction in a research study or clinical setting.

## Methods

To evaluate the different polygenic scoring approaches, we used two target samples: UK Biobank (UKB) [23], and the Twins Early Development Study (TEDS) [24]. All code used to prepare data and carryout analyses is available on the GenoPred website (see Data and Code Availability).

### UKB

UKB is a prospective cohort study that recruited >500,000 individuals aged between 40-69 years across the United Kingdom. The protocol and written consent were approved by the UKB’s Research Ethics Committee (Ref: 11/NW/0382).

#### Genetic data

UKB released imputed dosage data for 488,377 individuals and ∼96 million variants, generated using IMPUTE4 software [23] with the Haplotype Reference Consortium reference panel [25] and the UK10K Consortium reference panel [26]. This study retained individuals that were of European ancestry based on 4-means clustering on the first 2 principal components provided by the UKB (self-reported ancestry was not used), and removed related individuals (>3^rd^ degree relative) using relatedness kinship (KING) estimates provided by the UKB [23]. The imputed dosages were converted to hard-call format using a hard call threshold of zero.

#### Phenotype data

Nine UKB phenotypes were analyzed. Eight phenotypes were binary: Depression, Type II Diabetes (T2D), Coronary Artery Disease (CAD), Inflammatory Bowel Disorder (IBD), Rheumatoid arthritis (RheuArth), Multiple Sclerosis (MultiScler), Breast Cancer, and Prostate Cancer. Three phenotypes were continuous: Intelligence, Height, and Body Mass Index (BMI). Further information regarding outcome definitions can be found in the Supplementary Material.

Analysis was performed on a subset of ∼50,000 UKB participants for each outcome. For each continuous trait (Intelligence, Height, BMI), a random sample was selected. For disease traits, all cases were included, except for Depression and CAD where a random sample of 25,000 cases was selected. Controls were randomly selected to obtain a total sample size of 50,000. Sample sizes for each phenotype after genotype data quality control are shown in Table 1. Supplementary Figure S1 shows a schematic diagram of how UKB data was split into training and testing sample.

**Table 1.** Sample size of target sample phenotypes after quality control

| **UKB Phenotype** | **Description** | **Total sample size** | **No. of cases** | **No. of controls** |
| --- | --- | --- | --- | --- |
| Depression | Major depression | 50000 | 25000 | 25000 |
| Intelligence | Fluid intelligence | 50000 | NA | NA |
| BMI | Body Mass Index | 50000 | NA | NA |
| Height | Height | 50000 | NA | NA |
| T2D | Type-2 Diabetes | 50000 | 35112 | 14888 |
| CAD | Coronary Artery Disease | 50000 | 25000 | 25000 |
| IBD | Inflammatory Bowel Disease | 50000 | 46539 | 3461 |
| MultiScler | Multiple Sclerosis | 50000 | 48863 | 1137 |
| RheuArth | Rheumatoid Arthritis | 50000 | 46592 | 3408 |
| Prostate Cancer | Prostate Cancer | 50000 | 47073 | 2927 |
| Breast Cancer | Breast Cancer | 50000 | 41488 | 8512 |
| **TEDS Phenotype** |  |  |  |  |
| GCSE | Mean GCSE scores | 7296 | NA | NA |
| ADHD | ADHD symptoms | 7880 | NA | NA |
| BMI21 | Body Mass Index at age 21 | 5220 | NA | NA |
| Height21 | Height at age 21 | 5455 | NA | NA |

### TEDS

The Twins Early Development Study (TEDS) is a population-based longitudinal study of twins born in England and Wales between 1994 and 1996 [27]. Ethical approval for TEDS has been provided by the King’s College London ethics committee (reference: 05/Q0706/228). Written parental and/or self-consent was obtained before data collection. For this study, one individual from each twin pair was removed to retain only unrelated individuals.

#### Genetic data

TEDS participants were genotyped using two arrays, HumanOmniExpressExome-8v1.2 and AffymetrixGeneChip 6.0. Stringent quality control was performed separately for each array, prior to imputation via the Sanger Imputation server using the Haplotype Reference Consortium (release 1.1) reference data [25, 28]. Imputed genotype dosages were converted to hard-call format using a hard call threshold of 0.9, with variants for each individual set to missing if no genotype had a probability of >0.9. Variants with an INFO score < 0.4, MAF < 0.001, missingness > 0.05 or Hardy-Weinberg equilibrium p-value < 1×10^-6^ were removed.

#### Phenotypic data

This study used four continuous phenotypes within TEDS: Height, Body Mass Index (BMI), Educational Achievement, and Attention Deficit Hyperactivity Disorder (ADHD) symptom score (Table 1). These phenotypes were selected based on a previous polygenic study, enabling comparison across methods [29]. The phenotypes were derived using the same protocol as previously.

### Genotype-based Scoring

The following genotype-based scoring procedure provides reference standardized polygenic scores and can be applied to any datasets of imputed genome-wide array data (Figure 1).

**Figure 1.**
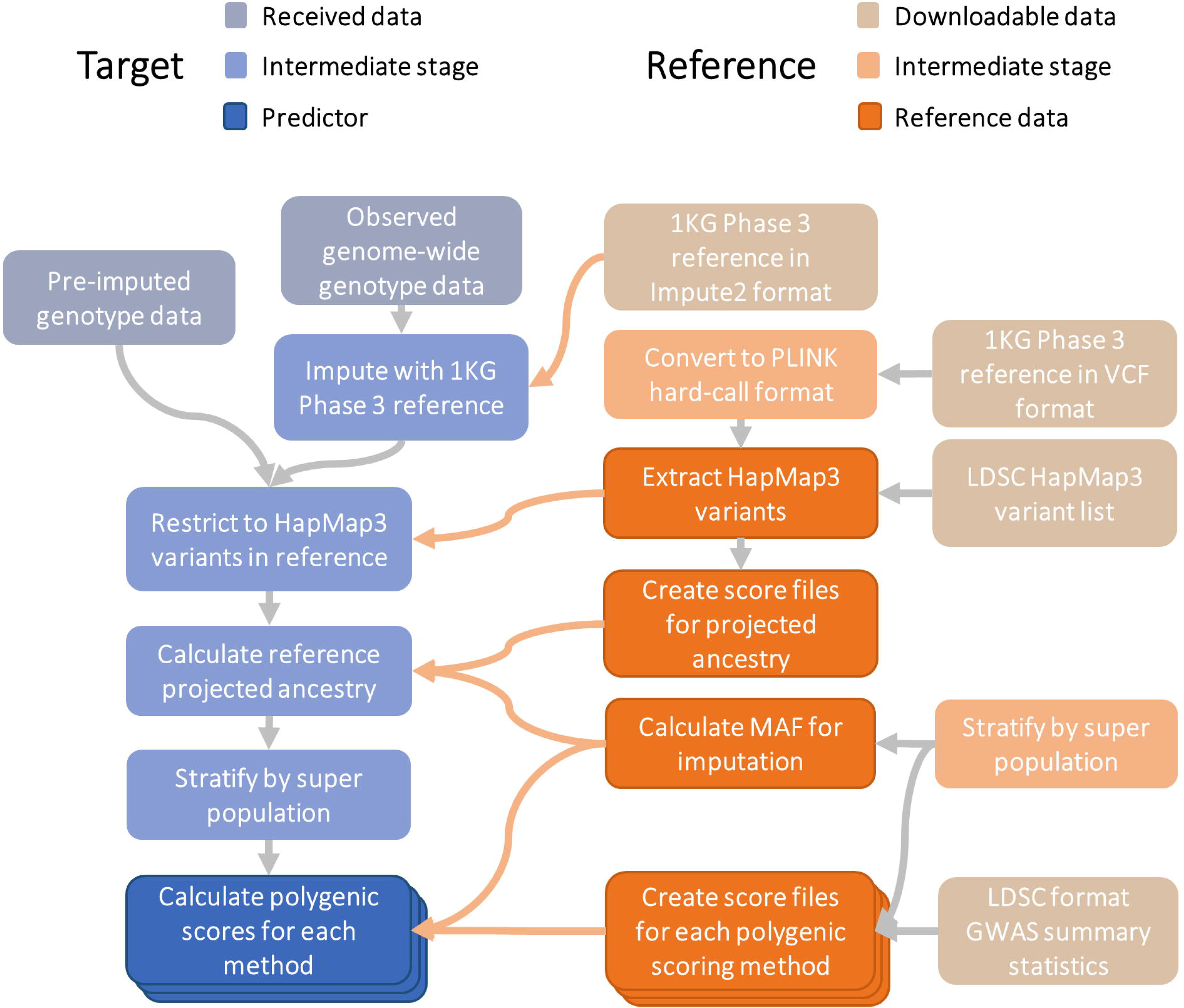
Schematic diagram of reference-standardized polygenic scoring. 1KG = 1000 Genomes; LDSC = Linkage Disequiibrium Score Regression; MAF = Minor allele Frequency; Pre-imputed genotype data = Indicates the observed genotype data has already been imputed; Observed genome-wide genotype data = Indicate the observed genotype data has not been imputed, and therefore requires imputation.

#### SNP-level QC

HapMap3 variants from the LD-score regression website (see Web Resources) were extracted from target samples (UKB, TEDS), inserting any HapMap3 variants that were not available in the target sample as missing genotypes (as required for reference MAF imputation by the PLINK allelic scoring function) [30]. No other SNP-level QC was performed.

#### Individual-level QC

Individual-level QC prior to imputation was previously performed for both UKB [23] and TEDS [28] samples. Only individuals of European ancestry were retained for polygenic score analysis. They were identified using 1000 Genomes Phase 3 projected principal components of population structure, retaining only those within three standard deviations from the mean for the top 100 principal components. This process will also remove individuals who are outliers due to technical genotyping or imputation errors.

#### GWAS summary statistics

GWAS summary statistics were identified for phenotypes the same as or similar as possible to the UKB and TEDS phenotypes (descriptive statistics in Table S1), excluding GWAS with documented sample overlap with the target samples. GWAS summary statistics underwent quality control to extract HapMap3 variants, remove ambiguous variants, remove variants with missing data, flip variants to match the reference, retain variants with a minor allele frequency (MAF) > 0.01 in the European subset of 1KG Phase 3, retain variants with a MAF > 0.01 in the GWAS sample (if available), retain variants with a INFO > 0.6 (if available), remove variants with a discordant MAF (>0.2) between the reference and GWAS sample (if available), remove variants with p-values >1 or </=0, remove duplicate variants, remove variants with sample size >3SD from the median sample size (if per variant sample size is available).

#### Reference genotype datasets

Target sample genotype-based scoring was performed using two different reference genotype datasets, the European subset of 1000 Genomes Phase 3 (N=503) and a random subset of 10,000 European-ancestry UKB participants. The UKB reference set was independent of the target sample used for evaluating polygenic scoring methods. These references were used to determine whether the sample size of the reference genotype dataset affects the prediction accuracy of polygenic scores. Only 1,042,377 HapMap3 variants were available in the UKB dataset and used in genotype-based scoring.

#### Polygenic Scores (PRS)

Polygenic scoring was carried out using eight approaches with default parameters outlined in Table 2. To ensure comparability across methods, the same set of HapMap3 variants were considered, and the same reference genotype datasets were used to estimate LD and MAF (except for PRScs and SBayesR).

**Table 2.** Description of polygenic scoring approaches.

| Method | Multiple tuning parameters | Pseudo-validation/ infinitesimal option | Software | Description | Parameters | MHC region | LD-reference |
| --- | --- | --- | --- | --- | --- | --- | --- |
| pT+clump[30] | Yes | No | PLINK | LD-based clumping and $p$-value thresholding | 10 nested $p$-value thresholds: 1e-8, 1e-6, 1e-4, 1e-2, 0.1, 0.2, 0.3, 0.4, 0.5, 1<br>Clumping: $r^2 = 0.1$; window = 250kb | Only top variant retained | EUR 1KG, EUR 10K UKB |
| lassosum[10] | Yes | Pseudo-validation | lassosum | Lasso regression-based | 80 $s$ and lambda combinations: $s = 0.2, 0.5, 0.9, 1$.<br>$\lambda = \exp(\text{seq}(\log(0.001), \log(0.1), \text{length.out}=20))^A$ | Not excluded | EUR 1KG, EUR 10K UKB |
| PRScs[11] | Yes | Pseudo-validation | PRScs | Bayesian shrinkage | 5 global shrinkage parameters ($\phi$) = 1e-6, 1e-4, 1e-2, 1, auto | Not excluded | PRScs-provided EUR 1KG |
| SBLUP[9] | No | Infinitesimal (only option) | GCTA | Best Linear Unbiased Prediction | NA | Not excluded | EUR 1KG, EUR 10K UKB |
| SBayesR[12] | No | Pseudo-validation (only option) | GCTB | Bayesian shrinkage | NA | Excluded (as recommended) | EUR 1KG, EUR 10K UKB, GCTB-provided |
| LDPred1[8] | Yes | Infinitesimal | LDPred | Bayesian shrinkage | Infinitesimal model and 7 non-zero effect fractions ($p$) = 3e-3, 1e-3, 3e-2, 1e-2, 3e-1, 1e-1, 1 | Not excluded | EUR 1KG, EUR 10K UKB |
| LDPred2[13] | Yes | Pseudo-validation and infinitesimal | bingsnpr | Bayesian Shrinkage | Auto, infinitesimal, and grid modes. Grid includes 126 combinations of heritability and non-zero effect fractions ($p$). | Not excluded | EUR 1KG, EUR 10K UKB |
| DBSLMM | No | Yes (only option) | DBSLMM | Bayesian shrinkage | NA | Not excluded | EUR 1KG, EUR 10K UKB |

*Note.* Default or recommended parameters were used for all methods.
<sup>A</sup>lassosum lambda values described using R code.

PRScs-provides an LD reference for HapMap3 variants based on the European subset of the 1000 Genomes, and results should be comparable to other methods when using the 1000 Genomes reference. PRScs was not applied using the larger UKB reference dataset as PRScs has been previously reported to show minimal improvement when using larger LD reference datasets [11].

SBayesR analysis requires shrunk and sparse LD matrices as input. LD matrices were calculated using Genome-wide Complex Trait Bayesian analysis (GCTB) [31] in batches of 5,000 variants, which were then merged for each chromosome, shrunk, and then made sparse. SBayesR analysis was also performed using LD matrices released by the developers of GCTB based on 50,000 European UKB individuals (see Web Resources).

Two additional modifications of the standard pT+clump approach were tested, termed ‘pT+clump (non-nested)’ and ‘pT+clump (dense)’. The pT+clump (non-nested) approach is the same the standard pT+clump approach except non-overlapping p-value thresholds were used to select variants included in the polygenic score, thereby making the polygenic scores for each threshold independent. The pT+clump (dense) approach is the same as the standard pT+clump approach except that it uses 10,000 p-value thresholds (minimum=5×10^-8^, maximum=0.5, interval=5×10-5), implemented using default settings in PRSice [7].

After adjustment of GWAS summary statistics as necessary for each polygenic scoring method, polygenic scores were calculated using PLINK with reference MAF imputation of missing data. All scores were standardized based on the mean and standard deviation of polygenic scores in the reference sample.

To determine whether certain methods are more prone to capturing genetic effects driven by population stratification, we carried out a sensitivity analysis, in which the first 20 principal components were regressed from the polygenic scores in advance. Principal components were derived in the 1KG Phase 3 reference, and then projected into UKB and TEDS samples.

#### Modelling approaches

For methods that provide polygenic scores based on a range of p-value thresholds (pT+clump) or shrinkage parameters (lassosum, PRScs, LDPred1, LDPred2), the best parameter was identified using either 10-fold cross validation (10FCVal) and, if available, pseudovalidation (PseudoVal). Pseudovalidation was performed using the pseudovalidate function in lassosum, the fully-Bayesian approach in PRScs, the auto model in LDPred2. SBayesR and DBSLMM by default estimate the optimal parameters and are therefore considered pseudovalidation methods. Methods assuming an infinitesimal model were SBLUP and the infinitesimal models of LDPred1 and 2. In addition to selecting the single ‘best’ parameter for polygenic scoring, elastic net models were derived containing polygenic scores based on a range of parameters for each method, with elastic net shrinkage parameters derived using 10-fold cross-validation (Multi-PRS). The number of scores generated by each method, which were included in the multi-PRS model, are shown in Table 2. In addition, we tested whether combining polygenic scores from all methods in an elastic net model improved prediction. This combined model is referred to the ‘All’ model.

The optimal parameters (pT, GWAS-effect size shrinkage, elastic net parameters) were determined based on the largest mean correlation between observed and predicted values obtained through 10-fold cross validation, and the resulting model was then applied to an independent test set. Ten-fold cross-validation is liable to overfitting when using penalized regression as hyperparameters are tuned using the 10-fold cross validation procedure. The independent test-set validation avoids any overfitting as the independent test sample is not used for hyperparameter tuning. Ten-fold cross validation was performed using 80% of the sample and the remaining 20% was used as the independent test sample. Ten-fold cross validation and test-set validation was carried out using the ‘caret’ R package, setting the same random seeds prior to subsetting individuals to ensure the same individuals were included for all polygenic scoring methods.

#### Evaluating prediction accuracy

Prediction accuracy was evaluated as the Pearson correlation between the observed and predicted outcome values. Correlation was used as the main test statistic as it is applicable for both binary and continuous outcomes and standard errors are easily computed as

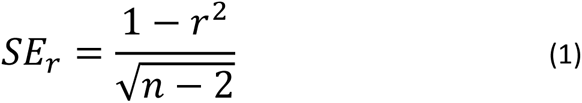

Where *SE_r_* is the standard error of the Pearson correlation, *r* is the Pearson correlation, and *n* is the sample size. Correlations can be easily converted to other test statistics such as *R^2^* (observed or liability) and area under the curve (AUC) (equations 8 and 11 in [32]), with relative performance of each method remaining unchanged.

When modelling the polygenic scores, logistic regression was used for predicting binary outcomes, and linear regression was used for predicting continuous outcomes. If the model contained only one predictor, a generalized linear model was used. If the model contained more than one predictor (i.e. the polygenic scores for each p-value threshold or shrinkage parameter), an elastic net model was applied to avoid overfitting due to the inclusion of multiple correlated predictors [33].

The correlation between observed and predicted values of each model were compared using William’s test (also known as the Hotelling-Williams test) [34] as implemented by the ‘psych’ R package’s ‘paired.r’ function, with the correlation between model predictions of each method specified to account for their non-independence. A two-sided test was used when calculating p-values.

The correlation between predicted and observed values were combined across phenotypes for each polygenic score method. Correlations and their variances (SE^2^) were aggregated using the ‘BHHR’ method [35] as implemented in the ‘MAd’ R package’s ‘agg’ function, using a phenotypic correlation matrix to account for the non-independence of analyses within each target sample. In addition to averaging results across all phenotypes, we estimate the average performance of methods within high and low polygenicity phenotypes. The polygenicity of phenotypes was estimated using AVENGEME [36] (more information in Supplementary Material).

The percentage difference between methods was calculated as

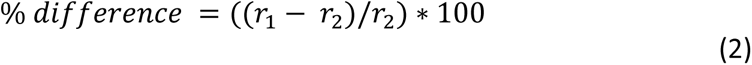

Where *r*_1_ and *r*_2_ indicate the Pearson correlation between predicted and observed values for models 1 and 2, respectively.

#### Method Runtime Comparison

To compare the time taken for each polygenic scoring method to process GWAS summary statistics, we ran each method using GWAS summary statistics restricted to variants on chromosome 22. No parallel implementations were used in this comparison. Apart from LDPred1, all the polygenic scoring methods can be implemented in parallel.

## Results

The eight polygenic risk score methods were applied to the target datasets of UKB (11 phenotypes) and TEDS (4 phenotypes), using two reference data sets of 1000 Genomes (1KG, 503 individuals) and UKB (10,000 individuals). Models were derived using 10-fold cross-validation, pseudovalidation, infinitesimal PRS and analysis of multiple threshold PRS, as appropriate for each polygenic risk score method (Table 2).

First, we confirmed that the design of the study was appropriate to detect differences between the methods using the GWAS summary statistics and test data sets chosen. GWAS summary statistics had sample sizes of a mean of 50,698 cases and 94,391 controls, and 423698 individuals for continuous traits, with heritability on the liability scale (estimated from the GWAS) ranging between 0.021 (Multiple Sclerosis) and 0.542 for Crohn’s disease (Table S1). For pT+clump, with 1KG reference and UKB target samples, the correlations between observed values (case-control status or measured trait) and the predicted values from the polygenic risk scoring models ranged from 0.074 (SE=0.010) for Intelligence to 0.299 (SE=0.010) for Height (Table S7). For each disorder or trait, reference panel and polygenic scoring method, the correlation was significantly different from zero (Tables S6-S9). These results confirm that the study design - comprising the GWAS, reference panel, target studies and traits - had sufficient information to capture polygenic prediction, and that the traits are diverse in polygenic architecture.

Results were highly concordant across the different target and reference samples used though the estimates were more precise when using the UKB target sample due to the increased sample size compared to TEDS (Figure S2-S3).

### Effect of reference panel and validation method

All polygenic scoring methods were applied to two reference panels of European ancestry: 503 individuals from the 1,000 Genomes sample, and 10,000 individuals from UKB. Results were highly similar for both panels (Figure S2-S3). For example, with the larger reference panel the correlation increased by a mean of 0.0017 in UKB, and 0.006 in TEDS, across traits and polygenic scoring methods (test-set validation, Table S2-S5; excluding PRScs which used only the 1,000 Genomes reference panel). Detailed results are reported here only for the 1,000 Genomes (1KG) reference panel, with full results for UKB reference panel in Supplementary Materials.

Both 10-fold cross validation and test-set validation methods were used in modelling, across all polygenic risk scoring methods. The 10-fold cross validation results were highly congruent with test-set validation results (Table 3). Results reported are based on test-set validation since this method is clearly robust to overfitting when using elastic net models (see Supplementary Materials for 10-fold cross-validation results).

**Table 3.** Average test-set correlation between predicted and observed values across phenotypes.

| Method | Model | CrossVal R (SE) | IndepVal R (SE) |
| --- | --- | --- | --- |
| pT+clump | 10FCVal | 0.155 (0.002) | 0.153 (0.004) |
| pT+clump | MultiPRS | 0.175 (0.002) | 0.174 (0.004) |
| lassosum | 10FCVal | 0.19 (0.002) | 0.183 (0.004) |
| lassosum | MultiPRS | 0.199 (0.002) | 0.194 (0.004) |
| lassosum | PseudoVal | 0.159 (0.002) | 0.157 (0.004) |
| PRScs | 10FCVal | 0.19 (0.002) | 0.183 (0.004) |
| PRScs | MultiPRS | 0.194 (0.002) | 0.187 (0.004) |
| PRScs | PseudoVal | 0.188 (0.002) | 0.182 (0.004) |
| SBLUP | Inf | 0.162 (0.002) | 0.156 (0.004) |
| SBayesR | PseudoVal | 0.097 (0.002) | 0.095 (0.004) |
| LDPred1 | 10FCVal | 0.178 (0.002) | 0.171 (0.004) |
| LDPred1 | MultiPRS | 0.181 (0.002) | 0.175 (0.004) |
| LDPred1 | Inf | 0.163 (0.002) | 0.156 (0.004) |
| LDPred2 | 10FCVal | 0.194 (0.002) | 0.187 (0.004) |
| LDPred2 | MultiPRS | 0.197 (0.002) | 0.191 (0.004) |
| LDPred2 | PseudoVal | 0.155 (0.002) | 0.151 (0.004) |
| LDPred2 | Inf | 0.161 (0.002) | 0.155 (0.004) |
| DBSLMM | PseudoVal | 0.182 (0.002) | 0.175 (0.004) |
| All | MultiPRS | 0.201 (0.002) | 0.196 (0.004) |

*Note.* This table shows results based on the UKB target sample and 1000 genomes reference. 10FCVal = Single polygenic score based on the optimal parameter as identified using 10-fold cross-validation. Multi-PRS = Elastic net model containing polygenic scores based on a range of parameters, with elastic net shrinkage parameters derived using 10-fold cross-validation. PseudoVal = Single polygenic score based on the predicted optimal parameter as identified using pseudovalidation, which requires no tuning sample, Inf = Single polygenic score based on infinitesimal model, which requires no tuning sample.

### Overview of polygenic scoring methods by modelling strategy

The performance for each polygenic scoring method across phenotypes was assessed using the correlation between observed and fitted values (Figure 2A), and then comparing each method with a baseline method of pT+clump with 10-fold cross validation using the difference in correlation (Figure 2B). All methods performed at least as well as pT+clump, except for SBayesR, which had convergence problems for several of the phenotypes (see Supplementary Material for full information). These overview results show that for the pseudovalidation (PseudoVal) and infinitesimal models (Inf) performed less well than polygenic scores selected through 10-fold cross-validation (10FCVal), and that the prediction when modelling multiple PRS (multi-PRS) was slightly higher than the 10-fold cross-validation. Full results for all traits in UKB and TEDS indicate consistency across methods, with no trait performing unexpectedly well or poorly on any single method (Tables S6-S9; Figures S4-S7).

**Figure 2.**
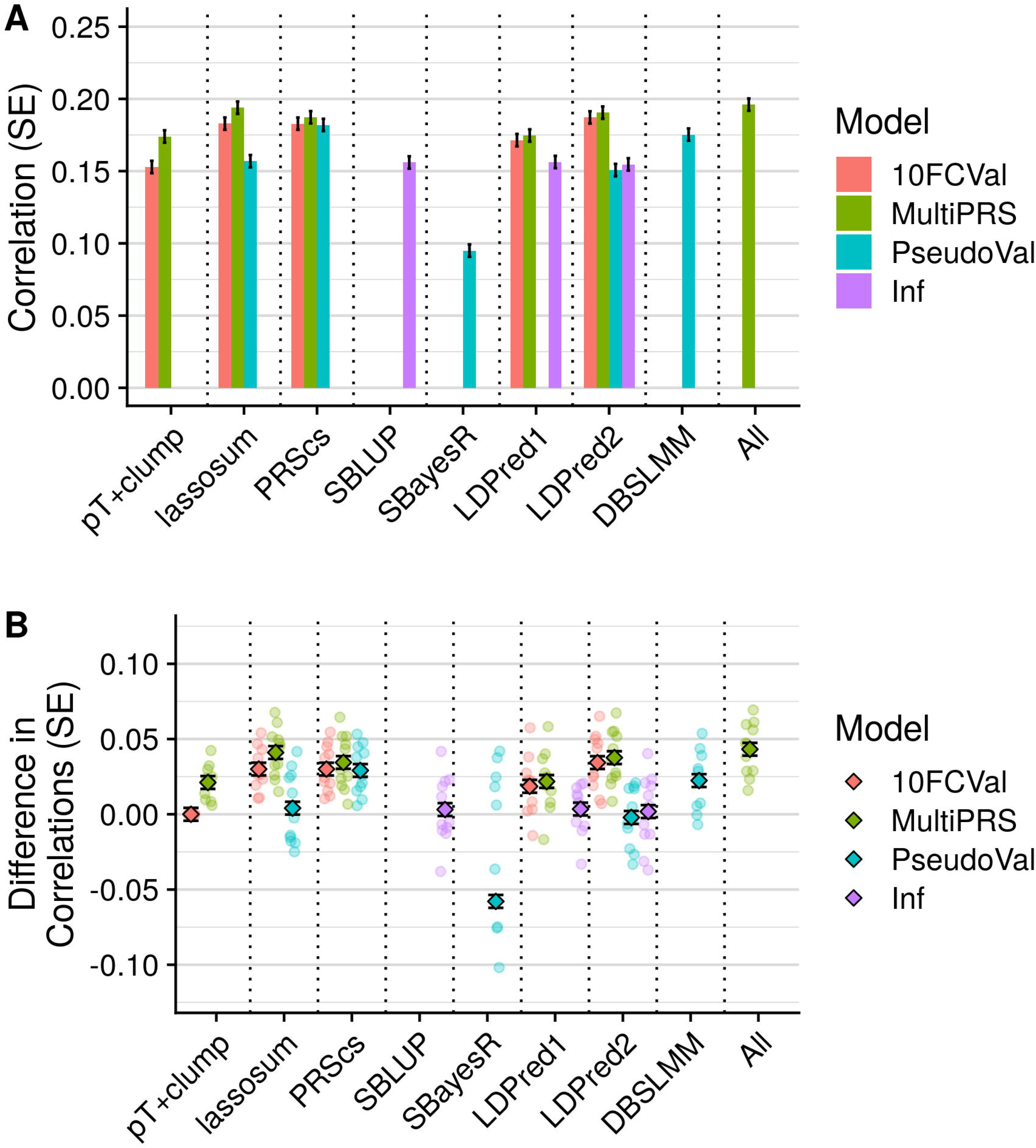
Polygenic scoring methods comparison for UKB target sample with 1KG reference. A) Average test-set correlation between predicted and observed values across phenotypes. B) Average difference between observed-prediction correlations for the best pT+clump polygenic score and all other methods. The average difference across phenotypes are shown as diamonds and the difference for each phenotype shown as transparent circles. SBayesR phenotype-specific correlation differences < −0.1 are omitted. Shows only results based on the UKB target sample when using the 1KG reference as other results were highly concordant. Error bars indicate standard error of correlations for each method. 10FCVal represents a single polygenic score based on the optimal parameter as identified using 10-fold cross-validation. Multi-PRS represents an elastic net model containing polygenic scores based on a range of parameters, with elastic net shrinkage parameters derived using 10-fold cross-validation. PseudoVal represents a single polygenic score based on the predicted optimal parameter as identified using pseudovalidation, which requires no tuning sample. Inf represents a single polygenic score based on the infinitesimal model, which requires no tuning sample.

### Comparison of polygenic scoring methods

A pairwise comparison of polygenic scoring methods was performed for each method (pT+clump, lassosum, PRScs, SBLUP, SBayesR, LDPred1, LDPred2, DBSLMM, All) and each model (10-fold cross validation, multi-PRS, pseudovalidation and infinitesimal). Figure 3 shows the difference in correlation (R) within and between methods for UKB outcomes with 1KG reference panel, with p-values for significant differences calculated using the William’s test results aggregated across outcomes. Full results for TEDS and UKB, and for both reference panels are given in Tables S10-S13 and Figure S8, and by trait in Tables S14-S17.

**Figure 3.**
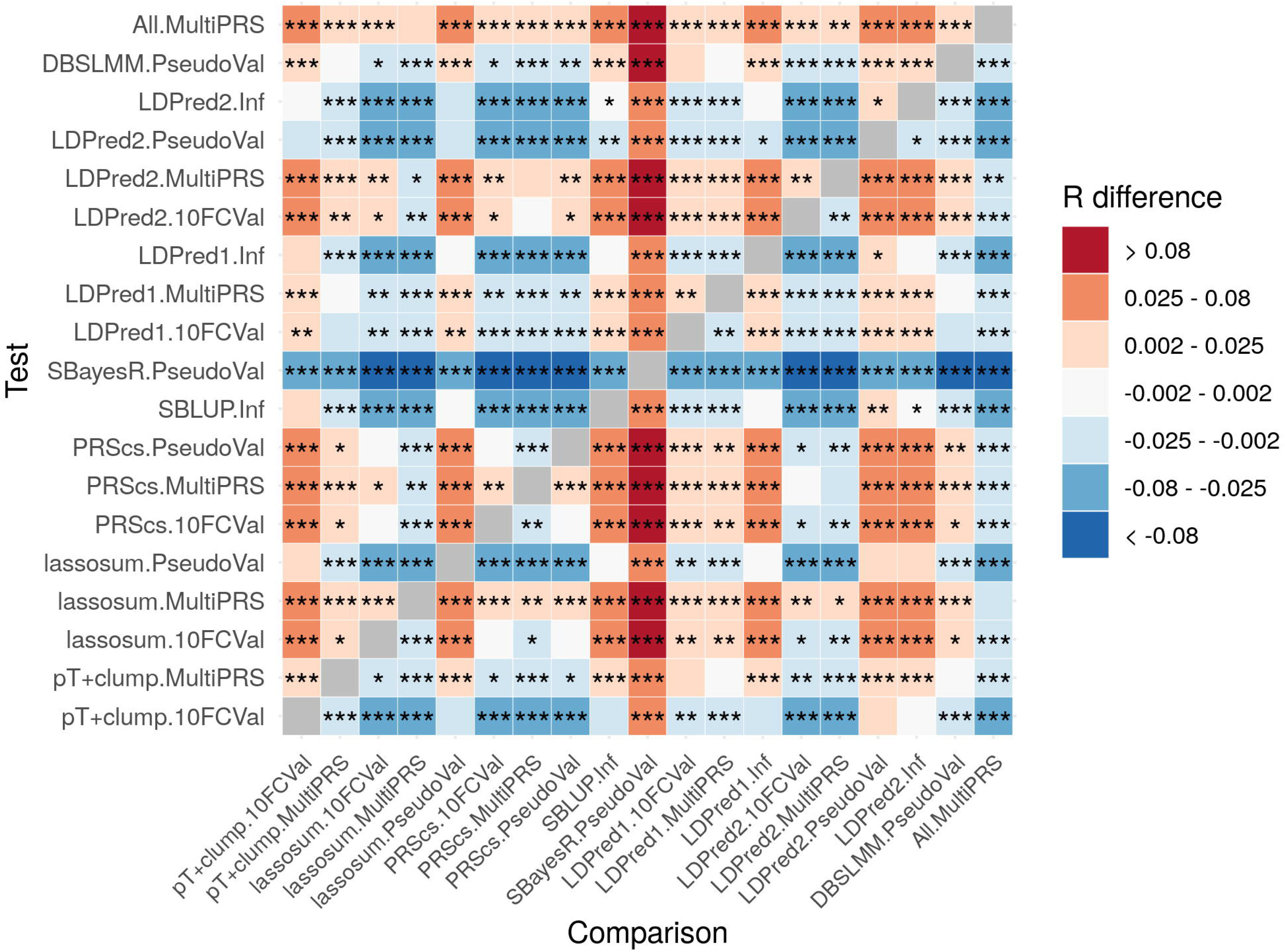
Pairwise comparison between all methods, showing average test-set observed-expected correlation difference between all methods with significance value. Correlation difference = Test correlation – Comparison correlation. Red/orange coloring indicates the Test method (shown on Y axis) performed better than the Comparison method (shown on X axis). Shows only results based on the UKB target sample when using the 1KG reference as other results were highly concordant. *=p<0.05. **=p<1×10^-3^. ***= p<1×10^-6^. P-values are two-sided. 10FCVal represents a single polygenic score based on the optimal parameter as identified using 10-fold cross-validation. Multi-PRS represents an elastic net model containing polygenic scores based on a range of parameters, with elastic net shrinkage parameters derived using 10-fold cross-validation. PseudoVal represents a single polygenic score based on the predicted optimal parameter as identified using pseudovalidation, which requires no tuning sample. Inf represents a single polygenic score based on the infinitesimal model, which requires no tuning sample.

When using 10-fold cross validation to identify the optimal parameter, LDPred2, lassosum and PRScs provided the most predictive polygenic scores in the test sample on average, with a 16-18% relative improvement (p<8×10^-16^) over the 10-fold cross-validated pT+clump approach. When using 10-fold cross validation, on average LDPred2 provided a small but nominally significantly improved prediction over lassosum and PRScs (2%, p=0.05).

Pseudovalidation and infinitesimal models do not require a tuning sample and their results are therefore described in parallel. Of the methods providing a pseudovalidation and/or infinitesimal approach (lassosum, PRScs, LDPred, LDPred2, SBLUP, DBSLMM and SBayesR), PRScs and DBSLMM performed the best on average, providing at least a 10% relative improvement (p<4×10^-9^) over other pseudovalidation approaches. The PRScs pseudovalidation approach provided a further significant improvement over DBSLMM, with an average relative improvement of 4% (p=4×10^-4^). Furthermore, the PRScs pseudovalidation approach was on average only 3% (*p*-value = 6×10^-3^) worse than the best polygenic score identified by 10-fold cross validation for any method. The performance of lassosum pseudovalidation, the LDPred1 and LDPred2 infinitesimal models, SBLUP, LDPred2 pseudovalidation and SBayesR was variable across phenotypes, whereas the PRScs pseudovalidated polygenic score achieved near optimal predication compared to any method, and always performed better than the best pT+clump polygenic scores as identified by 10-fold cross validation. The DBSLMM method performance was also relatively stable across phenotypes.

Modelling multiple polygenic scores based on multiple parameters using an elastic net consistently outperformed models containing the single best polygenic score as identified using 10-fold cross validation. The improvement was largest when using pT+clump polygenic scores (12% relative improvement, p=1×10^-21^), but was also statistically significant for lassosum (6% relative improvement, 3×10^-15^), PRScs (2% relative improvement, p=4×10^-5^), LDPred1 (2% relative improvement, p=4×10^-5^) and LDPred2 (2% relative improvement, p=3×10^-4^ methods. On average, the ‘All’ method, combining polygenic scores across polygenic scoring methods did not provide a statistically significant improvement over the single best method (multi-PRS lassosum). Elastic net models using non-nested or dense *p*-value thresholds showed no improvement over the standard *p*-value thresholding approach (Tables S18-S19).

The performance of SBayesR was higher when using the larger UKB reference sample (Figures S2-S3, S9-S10), though on average it still performed worse than all other approaches (Figure S8). SBayesR results based on the UKB reference were similar to those using the GCTB-provided LD reference (Figures S9-S10). The relative performance of SBayesR varied substantially (Figures 2B, S3, S9-S10). When using the UKB reference, the variable performance of SBayesR is partly due to a lack of convergence for Height, IBD and MultiScler, even when restricting variants to P<0.4 as suggested by the methods developers (Table S20). The SBayesR heritability results for each GWAS when using different approaches for preparing the summary statistics are shown in Table S20.

The relative performance of all methods and modelling approaches was similar across low and high polygenicity phenotypes (Figure S11). Infinitesimal model-based polygenic scores performed better for high polygenicity phenotypes. Estimates of polygenicity for each phenotype are shown in Table S21.

Controlling for the first 20 genetic principal components did not affect the relative performance of polygenic scoring methods (Figure S12).

### Runtime Comparison

The runtime of methods to process GWAS summary statistics on chromosome 22 without parallel implementations varied substantially (Figure S13). The methods (fastest to slowest) were pt+clump (∼3 seconds), DBSLMM and lassosum (∼30 seconds), SBLUP (∼1 minute), SBayesR and LDPred (∼3-6 minutes), PRScs (∼35 minutes), and LDPred2 (∼50 minutes). The number of parameters tested by each method will influence the runtime. For example, using only one shrinkage parameter for PRScs will take 1/5 of time taken for PRScs to use 5 shrinkage parameters.

## Discussion

This study evaluated a range of polygenic scoring methods across phenotypes representing a range of genetic architectures and using reference and target sample genotypic data of different sample sizes. This study shows that, when a tuning sample is available to identify optimal parameters, more recently developed methods that do not perform LD-based clumping provide better prediction, with LDPred2, lassosum and PRScs providing a relative improvement of 16-18% compared to the pT+clump approach. When a tuning sample is not available, the optimal methods for prediction was PRScs and DBSLMM, providing a >10% relative improvement over other pseudovalidation and infinitesimal approaches. The PRScs pseudovalidation method provided a further relative improvement of 4% over the DBSLMM method. Furthermore, the PRScs pseudovalidation performance was only 3% worse than the best polygenic scores identified by 10-fold cross validation for any other method. This study also shows that an elastic net model containing multiple polygenic scores based on a range of p-value thresholds or shrinkage parameters provides better prediction than the single best polygenic score as identified by 10-fold cross validation. Modelling multiple parameters increased prediction by 12% when using the pT+clump approach and 2-6% for polygenic scoring methods that model LD. Modelling polygenic scores from multiple methods did not significantly improve prediction over the single best method.

Our study highlighted the performance of SBayesR is highly variable across GWAS summary statistics and on average does not perform well compared to other methods. In contrast, a recent preprint comparing polygenic scoring methods using depression and schizophrenia GWAS reports that SBayesR is the best approach [16]. This apparent discrepancy can be explained by our study testing methods across a wider range of GWAS. Indeed, for the one phenotype tested in both studies (depression), the results are highly concordant, with SBayesR performing better than other methods when using larger LD-reference datasets. Our study highlights the importance of validating methods based on GWAS for a range of phenotypes and from different discovery samples/consortia.

These methods were evaluated within a reference-standardized framework and the results are likely to be generalizable to a range of settings, including a clinical setting. The improved transferability of prediction accuracy when using a reference-standardized approach enables prediction with a known accuracy for a single individual. This is an essential feature of any predictor as then its prediction can be appropriately considered in relation to other information about the individual. It is important to consider whether the reference-standardized approach impacts the predictive utility of the polygenic scores compared to those derived using target sample specific properties. The use of only HapMap3 variants is common for polygenic scoring methods as denser sets of variants increase the computational burden of the analysis and provide only incremental improvements in prediction [12]. However, denser sets of variants are ultimately likely to be of importance for optimizing the predictive utility of polygenic scores. The use of reference LD estimates instead of target sample-specific LD estimates is less likely to impact the predictive utility of polygenic scores. LD estimates are used to recapitulate LD structure in the GWAS discovery sample, and there should therefore be no advantage to using target sample specific LD estimates instead of reference sample LD estimates, unless the target sample better captures the LD structure in the GWAS discovery sample.

One major limitation of our study is that it was performed only in studies of European ancestry since GWAS of other ancestries have insufficient power for polygenic prediction. Polygenic scoring method comparisons in other ancestries or across ancestries will require substantial progress in diversifying genetic studies to non-European ancestry. In particular, it will be important to assess the impact of greater genetic diversity and weaker linkage disequilibrium in African ancestry populations. These studies are essential if polygenic risk scores are to be implemented in clinical care, to ensure equity of healthcare.

The clinical implementation of polygenic scores is at an early stage, and we identify five areas that still require further research. First, this study demonstrates that the reference-standardized approach provides reliable polygenic score estimates. However, the extent to which missing genetic variation within target sample data affects the prediction accuracy needs to be investigated. Furthermore, the extent to which prediction accuracy varies across individuals from different European ancestral populations needs to be assessed. Second, this study used the HapMap3 SNP list when deriving polygenic scores, building on previous research suggesting that these variants are reliably imputed and provide good coverage of the genome [20]. However, other sets of variants should be explored as denser coverage of the genome may improve prediction. Third, this study investigates polygenic scores based on a single discovery GWAS or phenotype. Previous research has shown that methods which combine evidence across multiple GWAS can improve prediction due to genetic correlation between traits [37–41]. Further research comparing the predictive utility of multi-trait polygenic prediction within a reference-standardized framework is required. Fourth, we present the reference standardized approach as a conceptual framework for implementing polygenic scores in a clinical setting. However, several additional issues will need to be addressed before they can be used in a clinical setting, such as assigning individuals to the optimal reference population, the presence of admixture, and translating relative polygenic scores into absolute terms. Finally, integration of functional genomic annotations has been shown to improve prediction over functionally agnostic polygenic scoring methods [42]. Comparison of functionally informed methods within a reference-standardized framework is also required.

In conclusion, this study performed a comprehensive comparison of GWAS summary statistic-based polygenic scoring methods within a reference-standardized framework using European ancestry studies. The results provide a useful resource for future research and endeavors to implement polygenic scores for individual-level prediction. All the code, rationale and results of this study are available on the GenoPred website (see Web Resources). This website will continue to document the evaluation of novel genotype-based prediction methods, providing a valuable community resource for education, research, and collaboration. Novel polygenic score methods can be rapidly tested against these standard methods to benchmark performance. This framework should be a valuable tool in the roadmap of moving polygenic risk scores from research studies to clinical implementation. Further investigation of methods providing genotype-based prediction within a reference-standardized framework is needed.

## Acknowledgements

We thank Luke Lloyd-Jones and Jian Zeng for their advice on SBayesR using GCTB, Tian Ge on using PRScs, and Florian Privé on using LDPred2. We thank Paul O’Reilly and Sam Choi for useful discussion.

This paper represents independent research funded by the UK Medical Research Council (MR/N015746/1 and MR/S0151132), and the National Institute for Health Research (NIHR) Biomedical Research Centre at South London and Maudsley NHS Foundation Trust and King’s College London. The authors acknowledge use of the research computing facility at King’s College London, Rosalind (https://rosalind.kcl.ac.uk), which is delivered in partnership with the NIHR Maudsley BRC, and part-funded by capital equipment grants from the Maudsley Charity (award 980) and Guy’s & St. Thomas’ Charity (TR130505). The views expressed are those of the authors and not necessarily those of the NHS, the NIHR or the Department of Health and Social Care. We thank the research participants and employees of 23andMe for making the work regarding Depression possible.

UKB: This research was conducted under UK Biobank application 18177.

TEDS: We gratefully acknowledge the ongoing contribution of the participants in TEDS and their families. TEDS is supported by UK Medical Research Council Program Grant MR/M021475/1 (and previously Grant G0901245) (to Robert Plomin (R.P).), with additional support from National Institutes of Health Grant AG046938. The research leading to these results has also received funding from the European Research Council under the European Union’s Seventh Framework Programme (FP7/2007-2013) ERC Grant Agreement 295366. R.P. is supported by Medical Research Council Research Professor Award G19/2. KR is supported by a Sir Henry Wellcome Postdoctoral Fellowship.

## Web Resources

- LDSC HapMap 3 SNP-list: https://data.broadinstitute.org/alkesgroup/LDSCORE/w_hm3.snplist.bz2
- LDSC Munge Sumstats: https://github.com/bulik/ldsc/blob/master/munge_sumstats.py
- GCTB LD matrices: https://zenodo.org/record/3350914
- Impute.me: https://impute.me/
- GenoPred: https://opain.github.io/GenoPred

## Data and Code Availability

The code used during this study are available at GitHub: https://opain.github.io/GenoPred.

An application is required to access individual-level data for TEDS and UKB.

