## Supplementary Information for "Evaluation of Polygenic Prediction Methodology within a Reference-Standardized Framework"

### UKB Outcome definitions

*Depression.* UKB participants were coded as depression cases if they met the Composite International Diagnostic Interview Short Form criteria for lifetime depression which was assessed in the online Mental Health Questionnaire (MHQ) using scoring protocols proposed by Davis et al [1]. Depression cases were screened for indications of schizophrenia or bipolar disorder according to the MHQ. Controls excluded if they show any psychiatric indications according to the MHQ or depression indications according to: ICD-10 diagnoses; endorsement of self-reported depression; endorsement of current antidepressant usage; single or current depression according to the criteria adopted by Smith, et al [2]. Further details of the exclusion criteria have been previously described [3].

*T2D.* Cases were identified based on a combination of hospital episode statistics, using both ICD-9 and ICD-10, the national death register, and self-reported questionnaire data. In order to classify as a case for type 2 diabetes, self-reported type 2 or generic diabetes status was established in the nurse interview and the touchscreen questionnaire. However, participants were only classified as cases when they reported in the questionnaire that they had not been treated with insulin in the first year after diagnosis and had been diagnosed after the age of 35 years. Type 2 diabetes controls did not fulfil these criteria and did not have any other types of diabetes. Further details of the T2D definition have been previously published [4].

*Coronary artery disease (CAD).* Participants who were registered in the hospital in-patient data or the death register to have had ischemic heart diseases, or participants who had coronary revascularization operations were classified as coronary artery disease cases in this study. If participants self-reported those conditions in the nurse interview or the touchscreen questionnaire, they were also considered to have coronary artery disease. Coronary artery disease controls did not fulfil those criteria. Further details of the CAD definition have been previously published [4].

*Autoimmune diseases (IBD, RheuArth, MultiScler):* UKB participants were coded as autoimmune cases if at least two of the following measures were observed: ICD-10 diagnoses from Hospital Episode Statistics; endorsement of self-reported autoimmune diseases; endorsement of prescription medication for the corresponding autoimmune diseases. More than one hospital admission for the respective autoimmune conditions was also sufficient. Controls were excluded if any of the following were observed: Pernicious Anemia, Autoimmune Thyroid Disease, Type 1 diabetes, Multiple Sclerosis, Myasthenia Gravis, Coeliac, Inflammatory Bowel Disease, Hidradenitis Suppurativa, Pemphigoid/Pemphigus, Psoriasis, Ankylosing Spondylitis, Polymyalgia Rheumatica/Giant Cell Arteritis, Psoriatic Arthritis, Rheumatoid Arthritis, Sjögren Syndrome, Systemic Lupus Erythematosus.

*Intelligence* was defined using the Fluid intelligence score variable. Fluid intelligence was assessed using the 13 item UKB Touch-screen Fluid intelligence test [5]. The test measures the capacity to solve problems that require logic and reasoning ability, independent of acquired knowledge. The fluid intelligence variable was derived by UKB as an unweighted sum of the number of correct answers, assigning a score of 0 to unanswered questions.

*Height* was defined using the Standing height variable (Field ID: f.50.0.0).

*BMI* was defined using the Body mass index variable (Field ID: f.21001.0.0).

Breast Cancer and Prostate Cancer were defined using the self-reported illness codes (1044 = prostate cancer, 1002 = breast cancer, Field ID: f.20001).

### TEDS outcome definitions

*Height*: Self-reported height was assessed at age 21.

*BMI*: BMI was calculated as self-reported weight in kilograms divided by height in meters squared (kg/m^2^) at age 21.

*Educational Achievement*: Results for standardized tests taken at the end of compulsory education in the United Kingdom (General Certificate of Secondary Education; GCSE) were obtained for twins at mean age 16.3 years (SD = 0.29) via self-report or parent-report. Grades were coded from 4 (G; the minimum pass grade) to 11 (A*; the highest possible grade), with the U fail grade coded as missing. A composite score was calculated as the arithmetic mean of the compulsory core subjects—Maths, English, and Science. Further information on this definition of Educational Attainment has been previously published [6].

*Attention-Deficit Hyperactivity Disorder (ADHD) Symptoms*: At age 11.5 (SD = 0.69) and 16.3 (SD = 0.69), parents reported on twins’ ADHD symptoms via the Strength and Difficulties Questionnaire [7] hyperactivity subscale (three-point Likert scale) and the Conners’ rating scales (CPRS-R; four-point Likert scale) [8] on hyperactivity and inattention. A composite score was created as the arithmetic mean of the sex and age z-standardized scales. Where ratings were available at one assessment only, this value was used to maximize sample size.

Estimating polygenicity

To investigate whether the polygenicity of the GWAS phenotype affects the relative performance of each polygenic scoring method, we averaged the predictive performance of each method across low polygenicity outcomes and high polygenicity outcomes. Polygenicity was estimated using software called AVENGEME [9], which uses pT+clump polygenic score association results across a range of p-value thresholds to estimate the polygenicity of the GWAS phenotype. We defined a GWAS phenotype as highly polygenic if the estimated proportion of variants with zero effect was <0.98. With this threshold the following outcomes were found to have low polygenicity: IBD, MultiScler, RheuArth, Prostate Cancer and Breast Cancer.

### SBayesR sensitivity analysis

SBayesR analysis can fail to converge due to its assumption that all variant effect sizes in the GWAS are estimated using the same number of individuals. A lack of convergence is indicated either by the SBayesR analysis not completing, or by a SNP-based heritability estimate of 1. To avoid convergence issues, we originally restricted the analysis to variants with a per-variant sample size within 3SD of the median sample size. However, for nine of the 12 GWAS, per variant sample size was not reported. As recommended by the SBayesR authors, for GWAS without per-variant sample size the GCTB ‘--impute-n’ option was used (imputes per variant sample size and removes variants with a sample size over 3SD from the median). Furthermore, if there was evidence of poor convergence, the analysis was restricted to variants with a p-value <0.4, as recommended by the SBayesR authors.

### Further SBayesR discussion

SBayesR has been previously reported to improve prediction over several of the methods tested in this study [10], yet within this study SBayesR performance varied substantially across phenotypes and on average performed worse than other methods. As previously reported, SBayesR is sensitive to the assumption that GWAS summary statistics are estimated using the same sample size for all variants, requiring truncation of the sample size variance (remove variants with sample sizes significantly different from the median). Truncating the sample size and restricting the analysis to variants with a GWAS p-value < 0.4 did improve the performance of SBayesR. However, the performance still varies substantially across phenotypes, on average performing worse than other methods. When using the UKB and GCTB-provided reference, this is partly due to unresolved convergence issues for 3 GWAS, including Height, IBD and MultiScler. Although SBayesR performed well for certain GWAS, further developments are required to improve the reliability of this method.

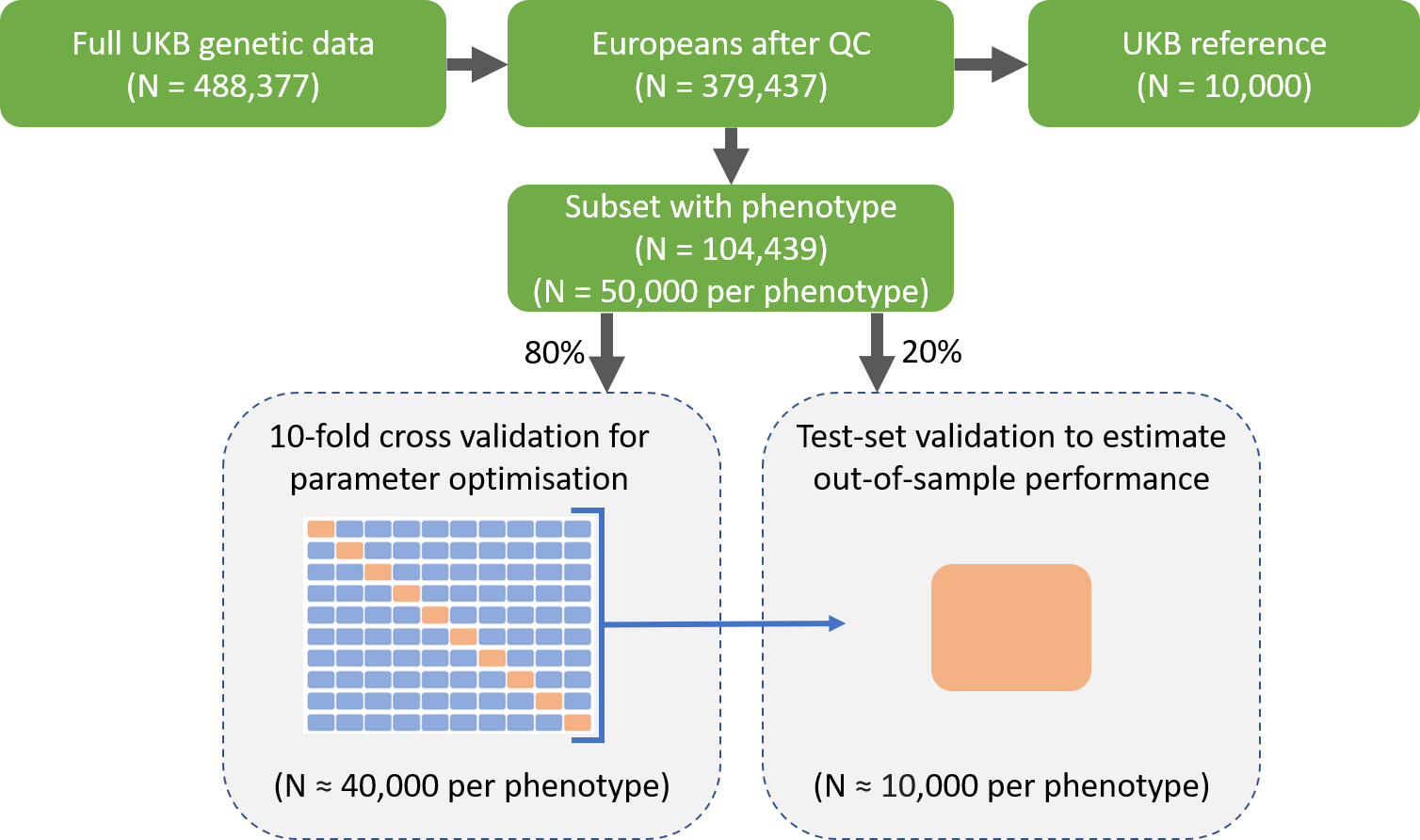

Supplementary Figure 1. Schematic diagram showing UKB was split into reference, training and testing samples. A sample of UKB providing 50,000 observations for each phenotype was identified. The sample was then further split into training (80%) and testing (20%) samples. The training sample used 10-fold cross validation to identify the optimal polygenic scoring parameters and elastic net hyper-parameters. An independent sample of 10,000 European UKB participants was also created to as a reference for polygenic scoring.

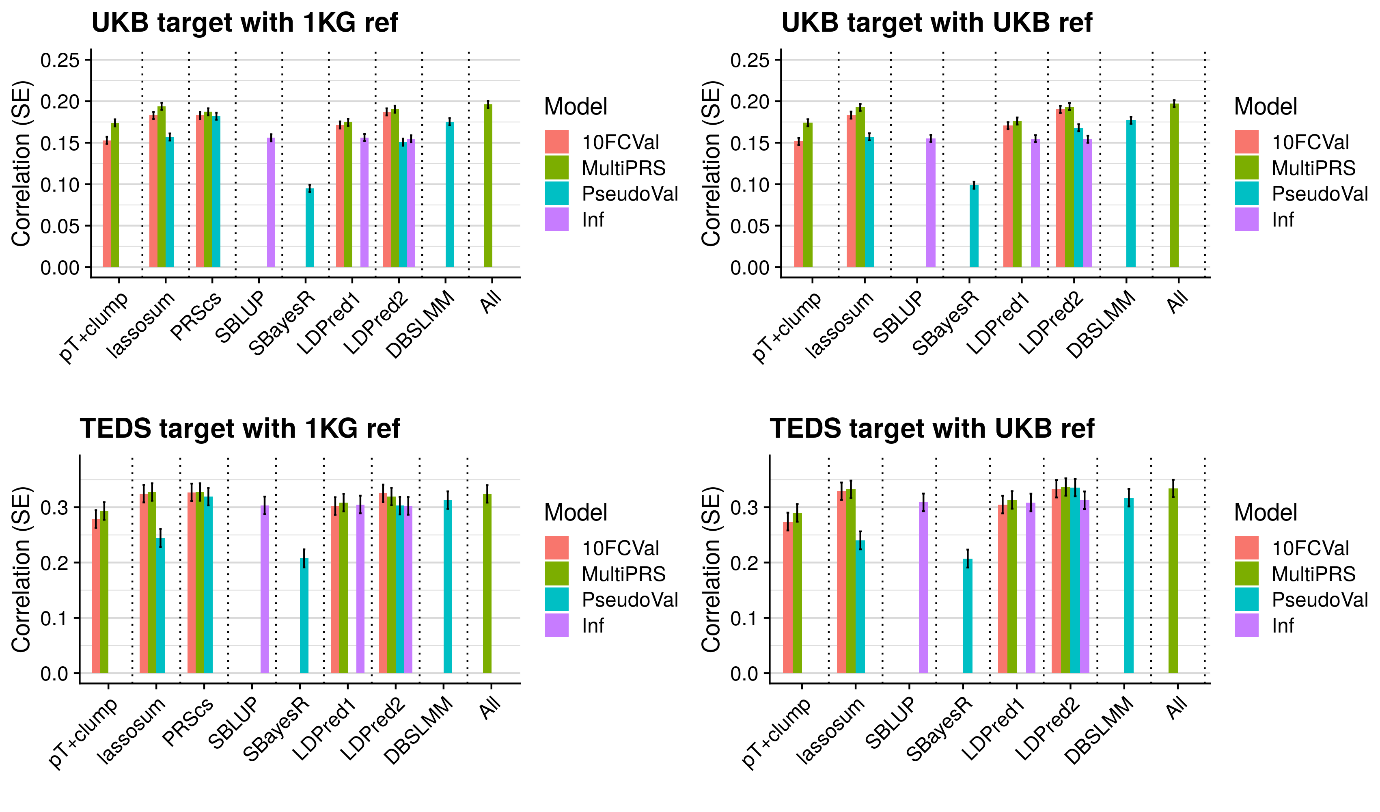

Supplementary Figure 2. Average test-set correlation between predicted and observed values across phenotypes. Error bars indicate standard error of correlations for each method. Results are split by the target and reference genotypic data used. Results are 10FCVal bars represent a single polygenic score based on the optimal parameter as identified using 10-fold cross-validation. Multi-PRS bars represent an elastic net model containing polygenic scores based on a range of parameters, with elastic net shrinkage parameters derived using 10-fold cross-validation. PseudoVal bars represent a single polygenic score based on the predicted optimal parameter as identified using pseudovalidation, which requires no tuning sample. Inf represents a single polygenic score based on the infinitesimal model, which requires no tuning sample.

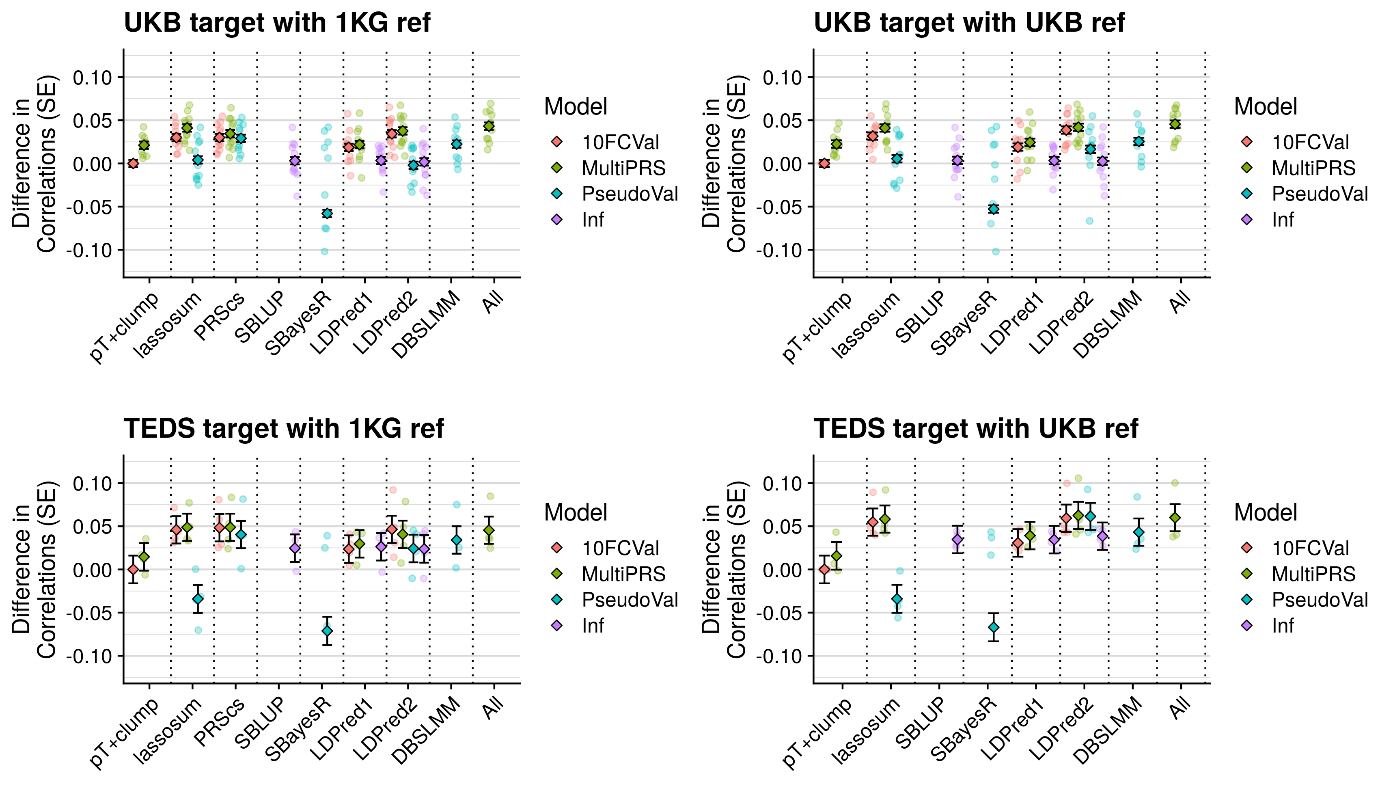

Supplementary Figure 3. Average test-set observed-expected correlation difference between the best pT+clump polygenic score and all other methods. The average difference across phenotypes are shown as diamonds with error bars indicating the standard error, and the difference for each phenotype shown as transparent circles. SBayesR phenotype-specific correlation differences < -0.1 are omitted. Results are split by the target and reference genotypic data used. 10FCVal represents a single polygenic score based on the optimal parameter as identified using 10-fold cross-validation. Multi-PRS represents an elastic net model containing polygenic scores based on a range of parameters, with elastic net shrinkage parameters derived using 10-fold cross-validation. PseudoVal represents a single polygenic score based on the predicted optimal parameter as identified using pseudovalidation, which requires no tuning sample. Inf represents a single polygenic score based on the infinitesimal model, which requires no tuning sample.

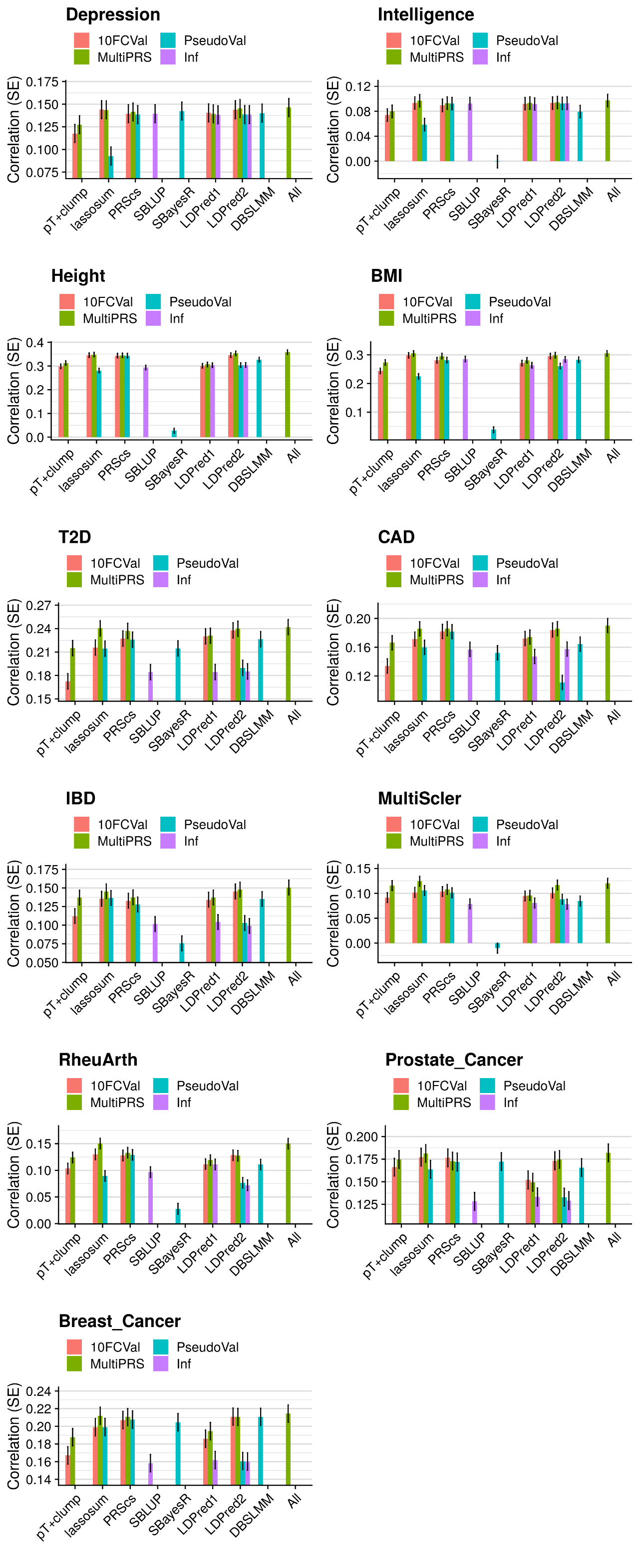

Supplementary Figure 4 Part 1. Correlation between predicted and observed values for each phenotype in UKB when using the European subset of 1000 Genomes as the reference. Error bars indicate standard errors. 10FCVal bars represent a single polygenic score based on the optimal parameter as identified using 10-fold cross-validation. Multi-PRS bars represent an elastic net model containing polygenic scores based on a range of parameters, with elastic net shrinkage parameters derived using 10-fold cross-validation. PseudoVal bars represent a single polygenic score based on the predicted optimal parameter as identified using pseudovalidation, which requires no tuning sample. Inf represents a single polygenic score based on the infinitesimal model, which requires no tuning sample.

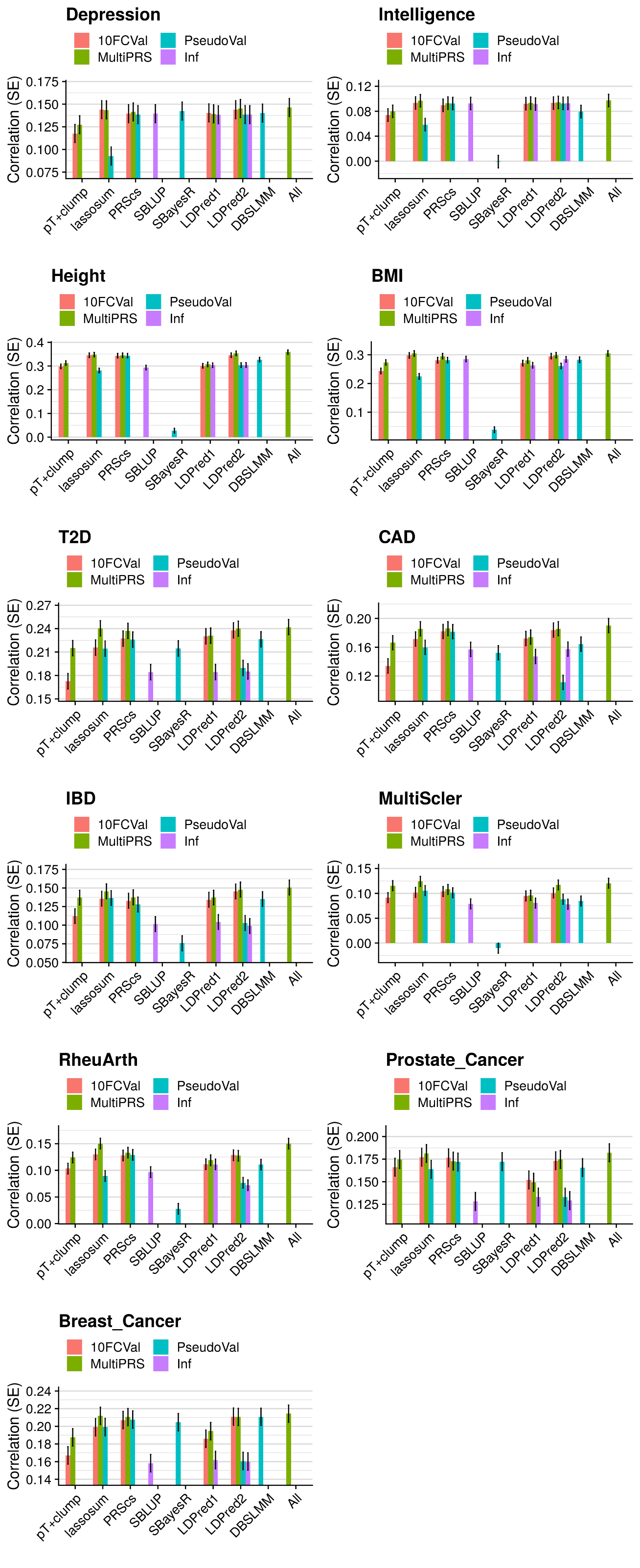

Supplementary Figure 4 Part 2. Correlation between predicted and observed values for each phenotype in UKB when using the European subset of 1000 Genomes as the reference. Error bars indicate standard errors. 10FCVal bars represent a single polygenic score based on the optimal parameter as identified using 10-fold cross-validation. Multi-PRS bars represent an elastic net model containing polygenic scores based on a range of parameters, with elastic net shrinkage parameters derived using 10-fold cross-validation. PseudoVal bars represent a single polygenic score based on the predicted optimal parameter as identified using pseudovalidation, which requires no tuning sample. Inf represents a single polygenic score based on the infinitesimal model, which requires no tuning sample.

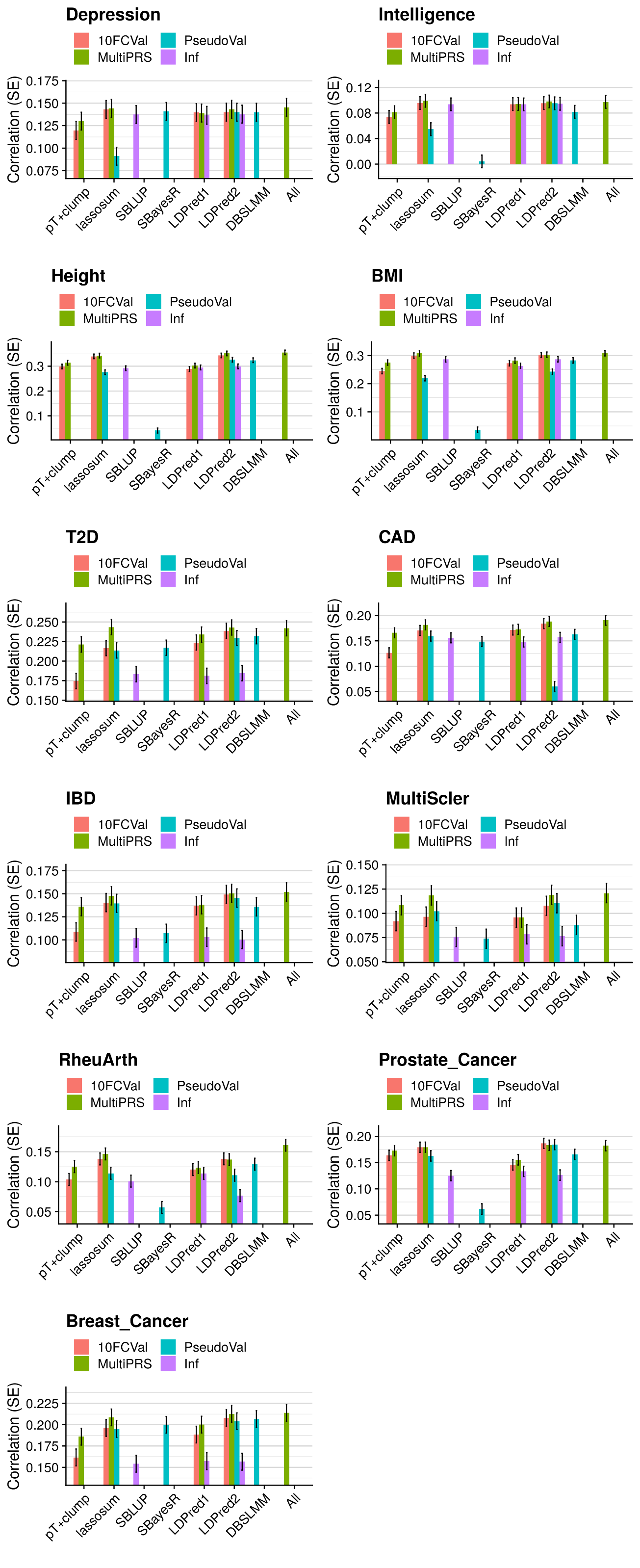

Supplementary Figure 5 Part 1. Correlation between predicted and observed values for each phenotype in UKB when using an independent 10K subset of European UKB individuals as the reference. Error bars indicate standard errors. 10FCVal bars represent a single polygenic score based on the optimal parameter as identified using 10-fold cross-validation. Multi-PRS bars represent an elastic net model containing polygenic scores based on a range of parameters, with elastic net shrinkage parameters derived using 10-fold cross-validation. PseudoVal bars represent a single polygenic score based on the predicted optimal parameter as identified using pseudovalidation, which requires no tuning sample. Inf represents a single polygenic score based on the infinitesimal model, which requires no tuning sample.

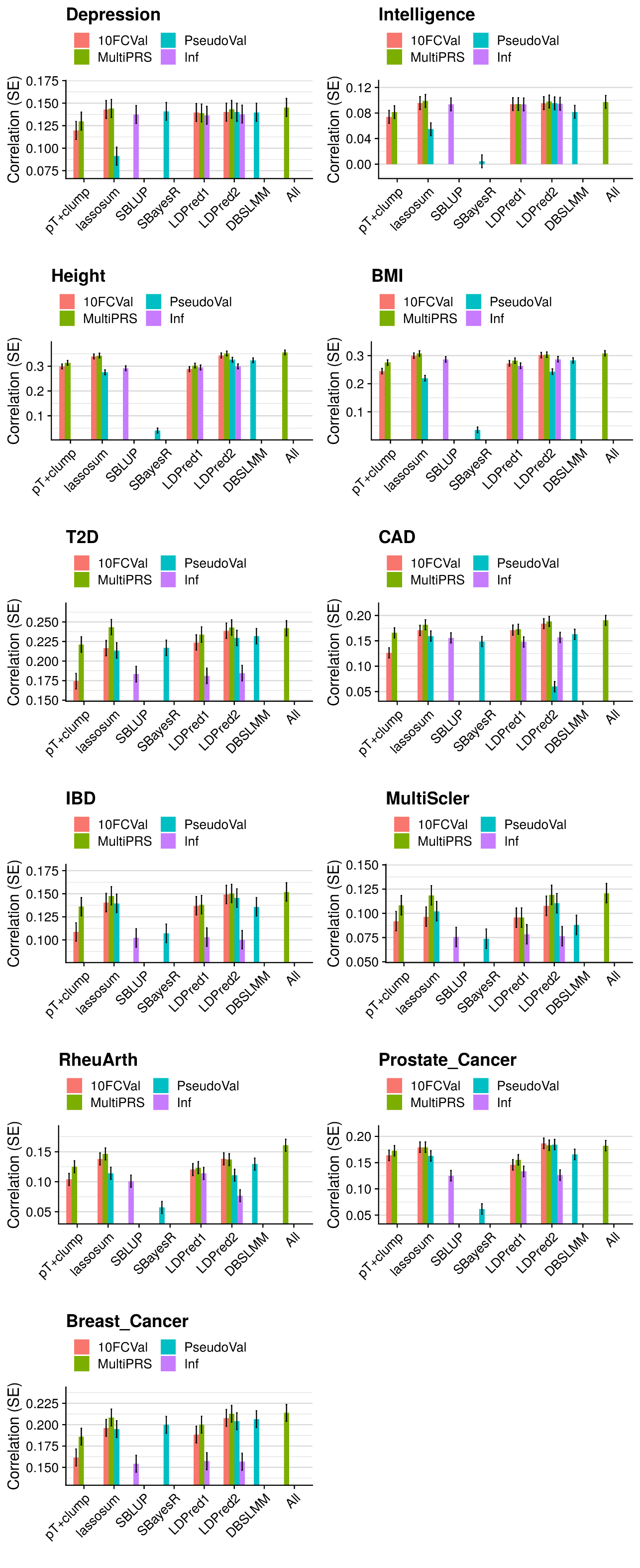

Supplementary Figure 5 Part 2. Correlation between predicted and observed values for each phenotype in UKB when using an independent 10K subset of European UKB individuals as the reference. Error bars indicate standard errors. 10FCVal bars represent a single polygenic score based on the optimal parameter as identified using 10-fold cross-validation. Multi-PRS bars represent an elastic net model containing polygenic scores based on a range of parameters, with elastic net shrinkage parameters derived using 10-fold cross-validation. PseudoVal bars represent a single polygenic score based on the predicted optimal parameter as identified using pseudovalidation, which requires no tuning sample. Inf represents a single polygenic score based on the infinitesimal model, which requires no tuning sample.

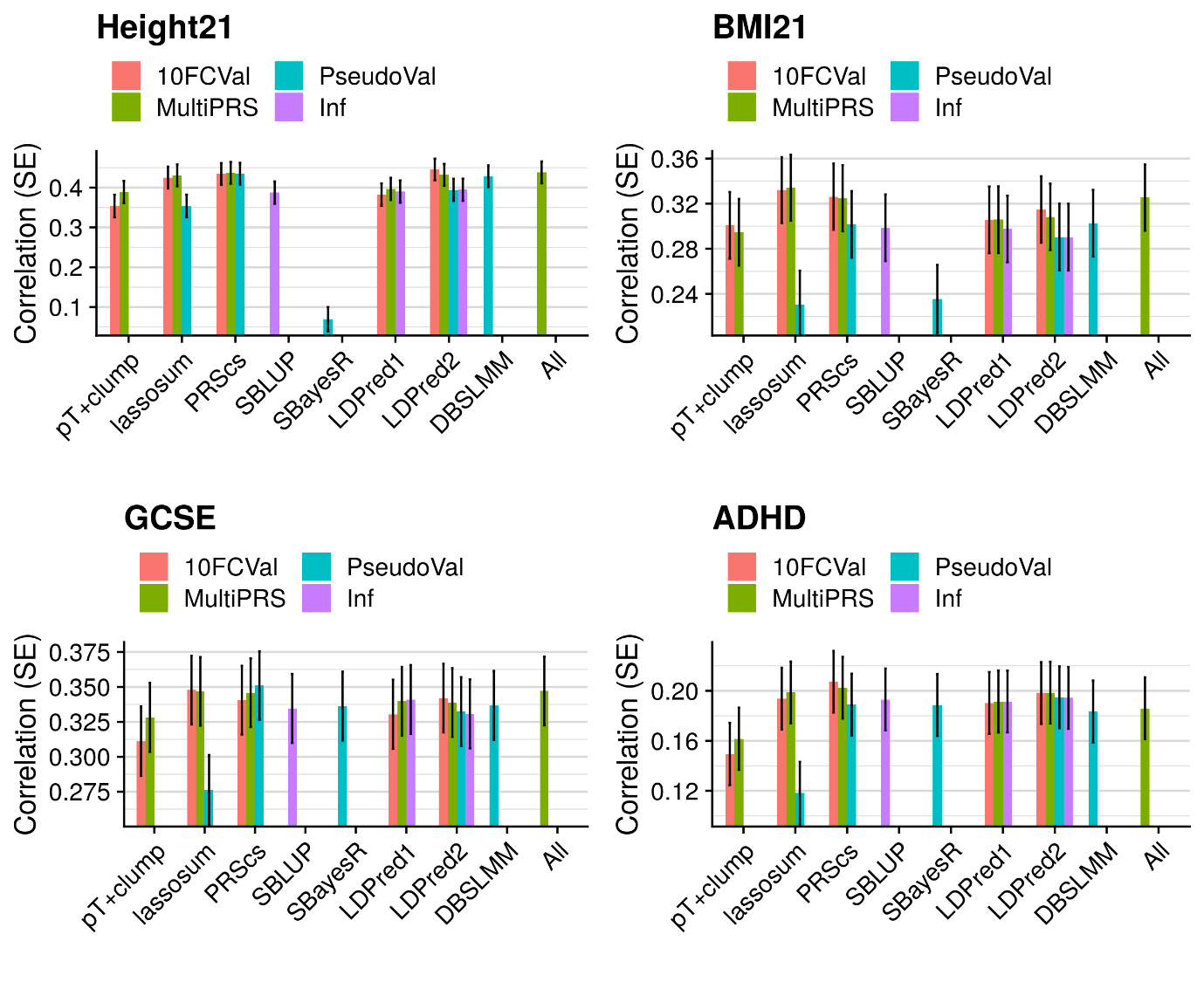

Supplementary Figure 6. Correlation between predicted and observed values for each phenotype in TEDS when using the European subset of 1000 Genomes as the reference. Error bars indicate standard errors. 10FCVal bars represent a single polygenic score based on the optimal parameter as identified using 10-fold cross-validation. Multi-PRS bars represent an elastic net model containing polygenic scores based on a range of parameters, with elastic net shrinkage parameters derived using 10-fold cross-validation. PseudoVal bars represent a single polygenic score based on the predicted optimal parameter as identified using pseudovalidation, which requires no tuning sample.1000G Reference. Inf represents a single polygenic score based on the infinitesimal model, which requires no tuning sample.

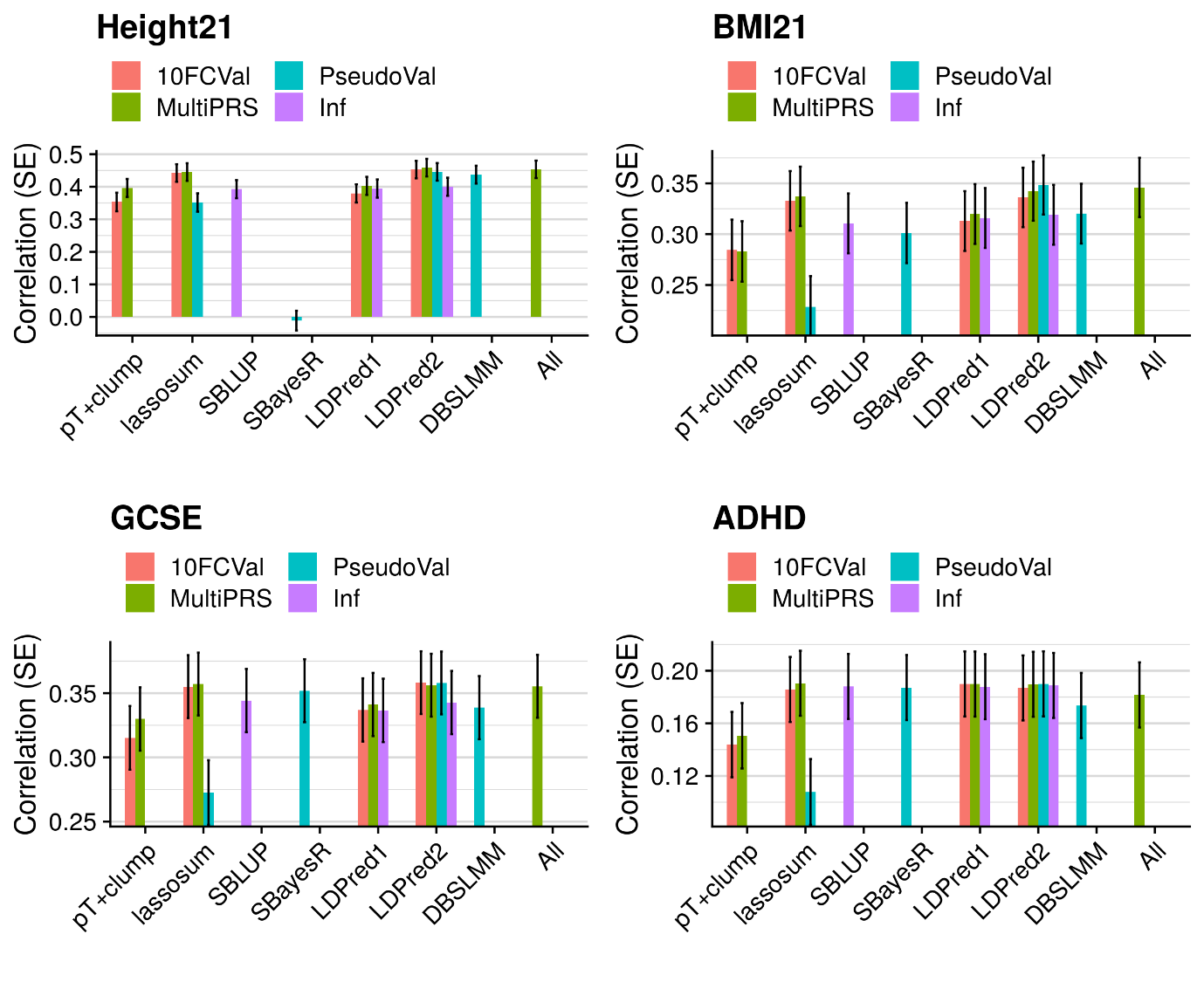

Supplementary Figure 7. Correlation between predicted and observed values for each phenotype in TEDS when using an independent 10K subset of European UKB individuals as the reference. Error bars indicate standard errors. 10FCVal bars represent a single polygenic score based on the optimal parameter as identified using 10-fold cross-validation. Multi-PRS bars represent an elastic net model containing polygenic scores based on a range of parameters, with elastic net shrinkage parameters derived using 10-fold cross-validation. PseudoVal bars represent a single polygenic score based on the predicted optimal parameter as identified using pseudovalidation, which requires no tuning sample. Inf represents a single polygenic score based on the infinitesimal model, which requires no tuning sample.

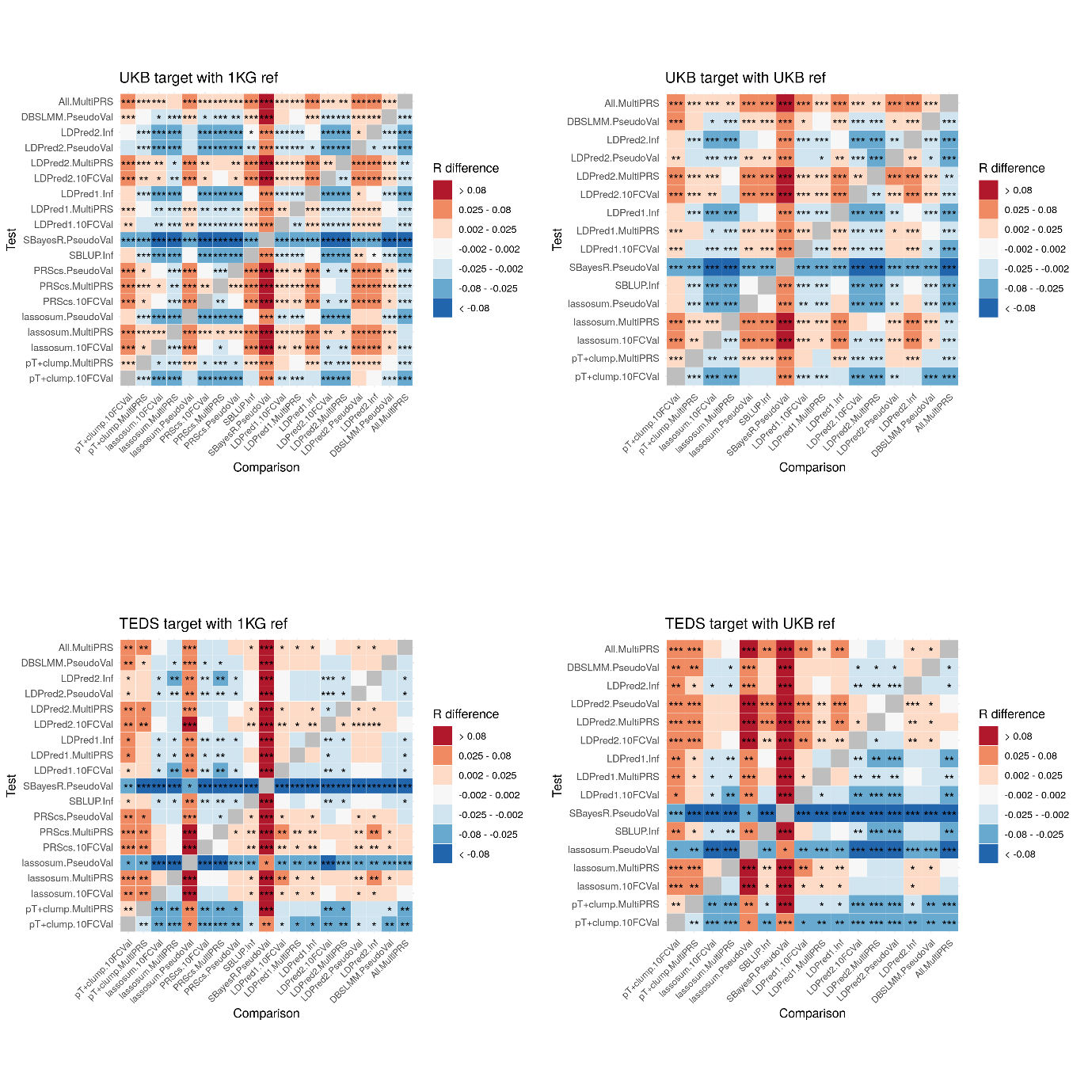

Supplementary Figure 8. Average test-set observed-expected correlation difference between all methods with significance value. Correlation difference = Test correlation – Reference correlation. Shows only results based on the UKB target sample when using the 1KG reference as other results were highly concordant. *=p<0.05. **=p<1×10^-3^. ***= p<1×10^-6^. P-values are one-sided. 10FCVal corresponds to a single polygenic score based on the optimal parameter as identified using 10-fold cross-validation. Multi-PRS corresponds to an elastic net model containing polygenic scores based on a range of parameters, with elastic net shrinkage parameters derived using 10-fold cross-validation. PseudoVal corresponds to a single polygenic score based on the predicted optimal parameter as identified using pseudovalidation, which requires no tuning sample. Inf represents a single polygenic score based on the infinitesimal model, which requires no tuning sample.

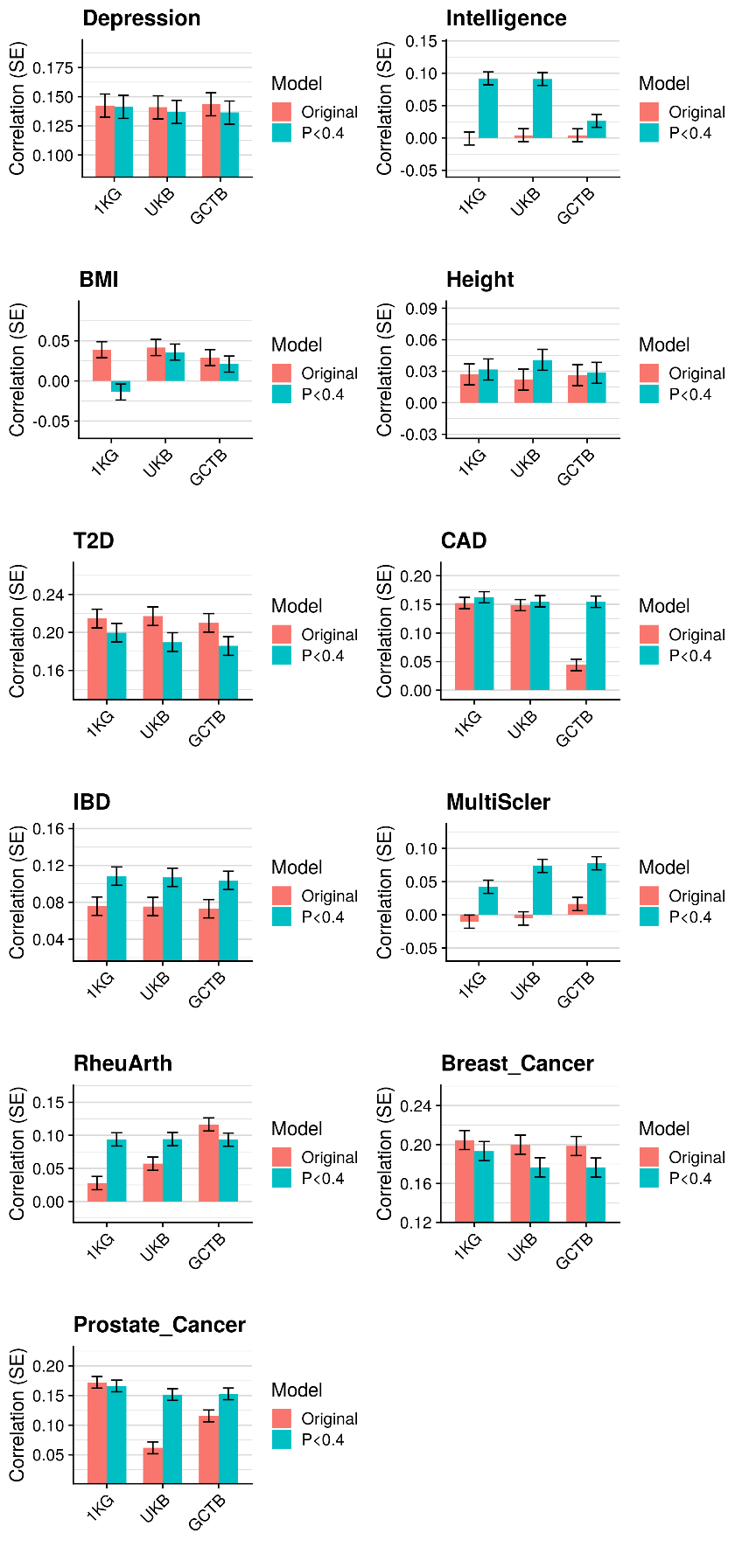

Supplementary Figure 9. Correlation between predicted and observed values across phenotypes in UKB for SBayesR polygenic scores derived using different reference samples and different GWAS processing procedures. Error bars indicate minimum and maximum correlations for each method. 1KG indicates the reference sample was the European subset of 1000 Genomes. UKB indicates the reference sample was an independent 10K subset of European UKB individuals. GCTB indicates the reference was the GCTB-provided reference data based on a non-independent 50K subset of European UKB individuals. Original indicates per variant sample size was imputed if missing and variants more than 3SD from the median sample size were removed. P<0.4 indicates only variants with a GWAS p-value <0.4 were retained.

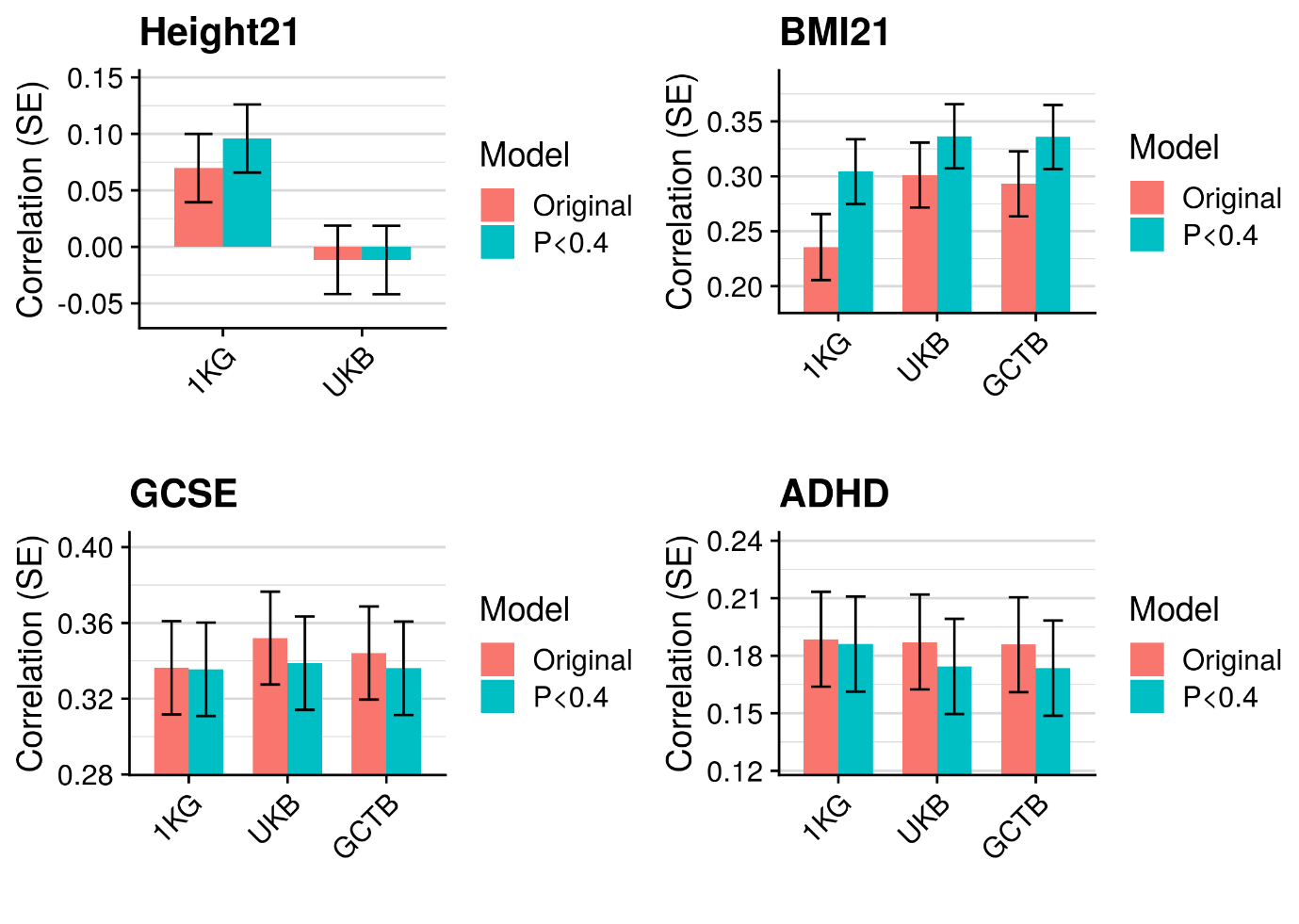

Supplementary Figure 10. Correlation between predicted and observed values across phenotypes in TEDS for SBayesR polygenic scores derived using different reference samples and different GWAS processing procedures. Error bars indicate minimum and maximum correlations for each method. 1KG indicates the reference sample was the European subset of 1000 Genomes. UKB indicates the reference sample was an independent 10K subset of European UKB individuals. GCTB indicates the reference was the GCTB-provided reference data based on a non-independent 50K subset of European UKB individuals. Original indicates per variant sample size was imputed if missing and variants more than 3SD from the median sample size were removed. P<0.4 indicates only variants with a GWAS p-value <0.4 were retained.

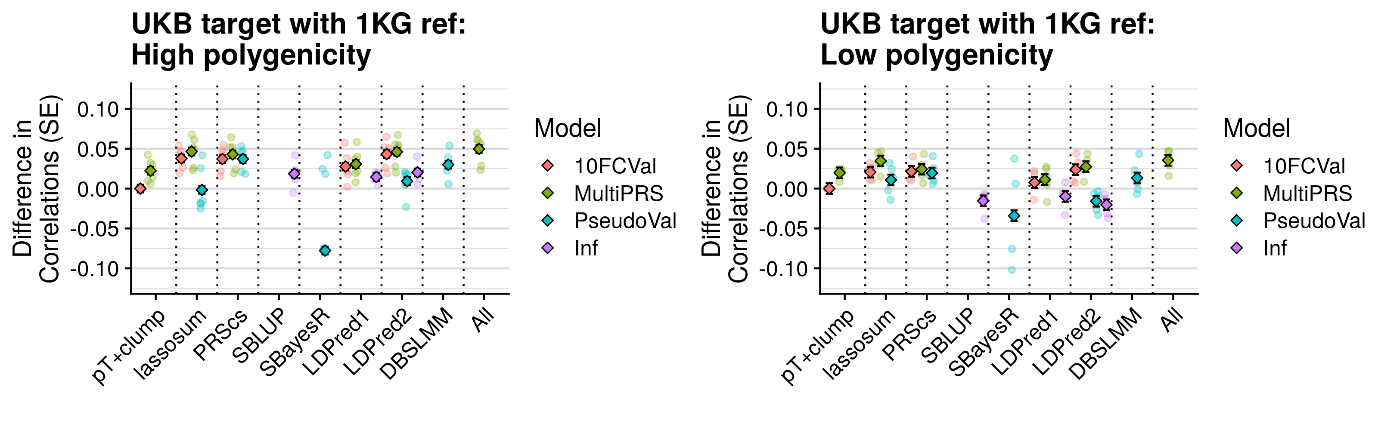

Supplementary Figure 11. Comparison of methods across high and low polygenicity outcomes in UKB target sample using 1KG reference. Figure shows average test-set observed-expected correlation difference between the best pT+clump polygenic score and all other methods. The average difference across phenotypes are shown as diamonds with error bars indicating the standard error, and the difference for each phenotype shown as transparent circles. SBayesR phenotype-specific correlation differences < -0.1 are omitted. 10FCVal represents a single polygenic score based on the optimal parameter as identified using 10-fold cross-validation. Multi-PRS represents an elastic net model containing polygenic scores based on a range of parameters, with elastic net shrinkage parameters derived using 10-fold cross-validation. PseudoVal represents a single polygenic score based on the predicted optimal parameter as identified using pseudovalidation, which requires no tuning sample. Inf represents a single polygenic score based on the infinitesimal model, which requires no tuning sample.

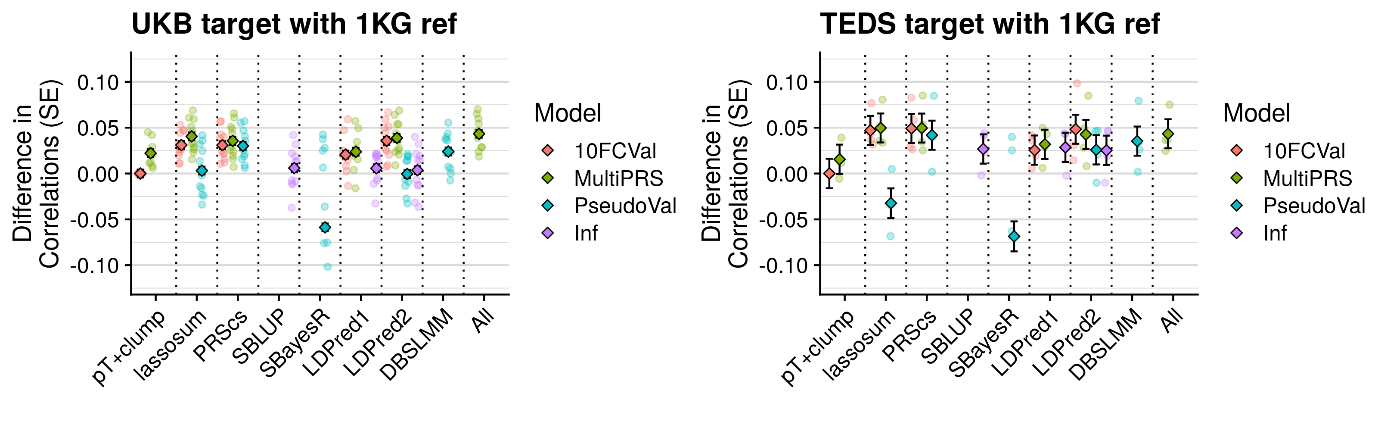

Supplementary Figure 12. Comparison of methods after controlling for genetic principal components in UKB target sample using 1KG reference. Figure shows average test-set observed-expected correlation difference between the best pT+clump polygenic score and all other methods. The average difference across phenotypes are shown as diamonds with error bars indicating the standard error, and the difference for each phenotype shown as transparent circles. SBayesR phenotype-specific correlation differences < -0.1 are omitted. 10FCVal represents a single polygenic score based on the optimal parameter as identified using 10-fold cross-validation. Multi-PRS represents an elastic net model containing polygenic scores based on a range of parameters, with elastic net shrinkage parameters derived using 10-fold cross-validation. PseudoVal represents a single polygenic score based on the predicted optimal parameter as identified using pseudovalidation, which requires no tuning sample. Inf represents a single polygenic score based on the infinitesimal model, which requires no tuning sample.

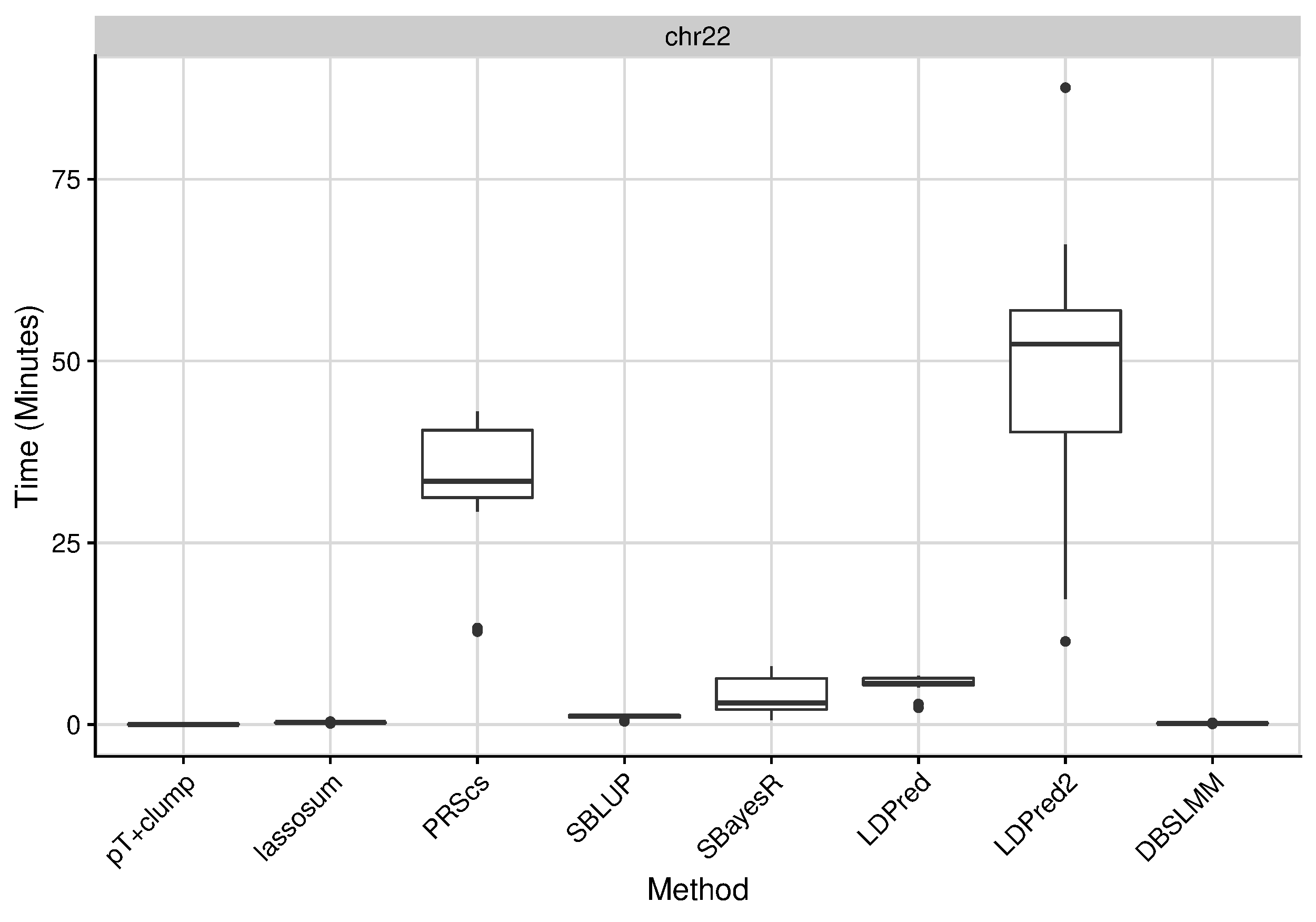

Supplementary Figure 13. Runtime for each polygenic scoring method using genetic variants on chromosome 22. No parallel implementations were used.
